# Intranasal self-amplifying RNA SARS-CoV-2 vaccine produces protective respiratory and systemic immunity and prevents viral transmission

**DOI:** 10.1101/2022.11.10.515993

**Authors:** Madeleine F. Jennewein, Michael D. Schultz, Samuel Beaver, Peter Battisti, Julie Bakken, Derek Hanson, Jobaida Akther, Fen Zhou, Raodoh Mohamath, Jasneet Singh, Noah Cross, Darshan N. Kasal, Matthew Ykema, Sierra Reed, Davies Kalange, Isabella R. Cheatwood, Jennifer L. Tipper, Jeremy B. Foote, R. Glenn King, Aaron Silva-Sanchez, Kevin S. Harrod, Davide Botta, Alana Gerhardt, Corey Casper, Troy D. Randall, Frances E. Lund, Emily A. Voigt

## Abstract

While mRNA vaccines have been effective in combating SARS-CoV-2, waning of vaccine-induced antibody responses and lack of vaccine-induced respiratory tract immunity contribute to ongoing infection and transmission. In this work, we compare and contrast intranasal (i.n.) and intramuscular (i.m.) administration of a SARS-CoV-2 self-amplifying RNA (saRNA) vaccine delivered by a nanostructured lipid carrier (NLC). Both i.m. and i.n. vaccines induce potent systemic serum neutralizing antibodies, bone marrow-resident IgG-secreting cells, and splenic T cell responses. The i.n. vaccine additionally induces robust respiratory mucosal immune responses, including SARS-CoV-2-reactive lung-resident memory and lung-homing T cell populations. As a booster following previous i.m. vaccination, the i.n. vaccine also elicits the development of mucosal virus-specific T cells. Both the i.m. and i.n. administered vaccines durably protect hamsters from infection-associated morbidity upon viral challenge, significantly reducing viral loads and preventing challenged hamsters from transmitting virus to naïve cagemates. This saRNA-NLC vaccine’s potent systemic immunogenicity, and additional mucosal immunogenicity when delivered i.n., may be key for combating SARS-CoV-2 and other respiratory pathogens.

## INTRODUCTION

The severe acute respiratory syndrome coronavirus 2 (SARS-CoV-2) pandemic prompted the rapid development and advancement of novel vaccines for coronavirus disease 2019 (COVID-19). Since their introduction in late 2020, two now FDA-approved mRNA vaccines (Comirnaty and Spikevax) have had a remarkable impact on the trajectory of the pandemic and were developed at unprecedented speeds ^1^. However, while these vaccines displayed outstanding efficacy at preventing severe COVID-19 in immunocompetent individuals and in reducing global mortality, the absence of robust vaccine-induced local immunity in the respiratory tract allows for persistent viral replication at the nasal mucosa in vaccinated individuals and subsequent viral transmission ^2–4^. In addition to the current COVID mRNA vaccines’ limited mucosal immunity, other challenges facing current vaccines include constantly evolving viral variants ^5^, rapidly waning efficacy ^6^ necessitating frequent booster doses ^7^ with diminishing uptake, vaccine hesitancy in part fueled by a fear of needles ^8^, and limited access to cold chain storage ^9^; global susceptibility to SARS-CoV-2 remains high. Improved vaccines and associated technologies are therefore necessary to fully combat endemic spread of the virus and, most importantly, effectively combat potential future respiratory virus pandemics.

We previously developed an intramuscularly (i.m.) administered nanostructured lipid carrier (NLC)-delivered self-amplifying RNA (saRNA) vaccine against SARS-CoV-2 ^10,11^, which induces both high serum neutralizing responses ^12–14^ and robust activation of systemic antigen-specific CD4^+^ and CD8^+^ T cells, responses that may be critical to sustained and potent immunity^13^. This saRNA-NLC vaccine platform additionally has unique thermostability relative to other mRNA delivery systems ^11,15–17^, as it may be readily lyophilized and stored at room temperature for months or under refrigeration likely for years ^11,17^.

We wanted to examine the potential for this vaccine, when administered intranasally (i.n.), to stimulate mucosal immune responses ^18^, which may be key to improving sustained, potent immunity to SARS-CoV-2 and other respiratory pathogens. Unlike i.m. administered vaccines, i.n. administered vaccines can induce critical immune cells within the mucosa and respiratory tract, generating IgA-secreting cells as well as specific effector cell subsets, including tissue-resident memory (T_RM_) T cells that are maintained within the lung ^19–26^. Such mucosal immunity, mediated by IgA antibodies and lung-resident memory T and B cells ^27^, can prevent early-stage infection and viral replication and shedding, minimizing chances for viral transmission ^20,21,26,28,29^. Effective i.n. administered vaccines can also induce more cross-reactive IgA antibodies and thus provide expanded protection to multiple strains ^30^. Intranasal vaccination also presents many other advantages, such as easier needle-free administration, higher patient compliance, simpler pediatric dosing, and easier storage and transport ^31^, together potentially increasing vaccine uptake. However, while several i.n. vaccines are currently under development for SARS-CoV-2, most rely on older vaccine technologies, i.e., live-attenuated viruses and viral vectors ^1^, creating a slow pathway to manufacturing, testing, and licensing. Fortunately, bridging RNA vaccine technologies to i.n. delivery is possible, combining the production speed of RNA vaccines with the mucosal immune stimulation of i.n. vaccines.

In this work, we demonstrate that both i.n. and i.m. administrations of our saRNA SARS-CoV-2 vaccine strongly induce systemic immunity in preclinical models and prevent viral transmission by vaccinated, virally-challenged hamsters. Intranasal dosing uniquely induces strong mucosal T cell immunity, and a heterologous dosing strategy—i.m. prime followed by i.n. boosting— appears to show the greatest potential to maximize mucosal and systemic immunity, leading to prevention of viral transmission. This saRNA-NLC vaccine candidate combines the flexibility and manufacturability of RNA vaccines with potent induction of mucosal immunity using a thermostable formulation that represents a significant new tool for current and future pandemic responses.

## RESULTS

### Intranasal saRNA-NLC vaccine stimulates robust systemic immune responses

For development of an intranasally delivered saRNA vaccine against SARS-CoV-2, we harnessed our previously developed and published long-term thermostable SARS-CoV-2 saRNA-NLC vaccine ^11^. This vaccine, previously tested in a Phase 1 clinical trial using i.m. administered material ^32^, contains a codon-optimized Wuhan-strain D614G SARS-CoV-2 spike protein sequence driven by a Venezuelan equine encephalitis virus (VEEV)-based replicon (Figure 1A) ^11^. This saRNA is complexed to the exterior of AAHI’s proprietary NLC nanoparticle to create the final vaccine product AAHI-SC2 (Figure 1B).

**Fig. 1.**
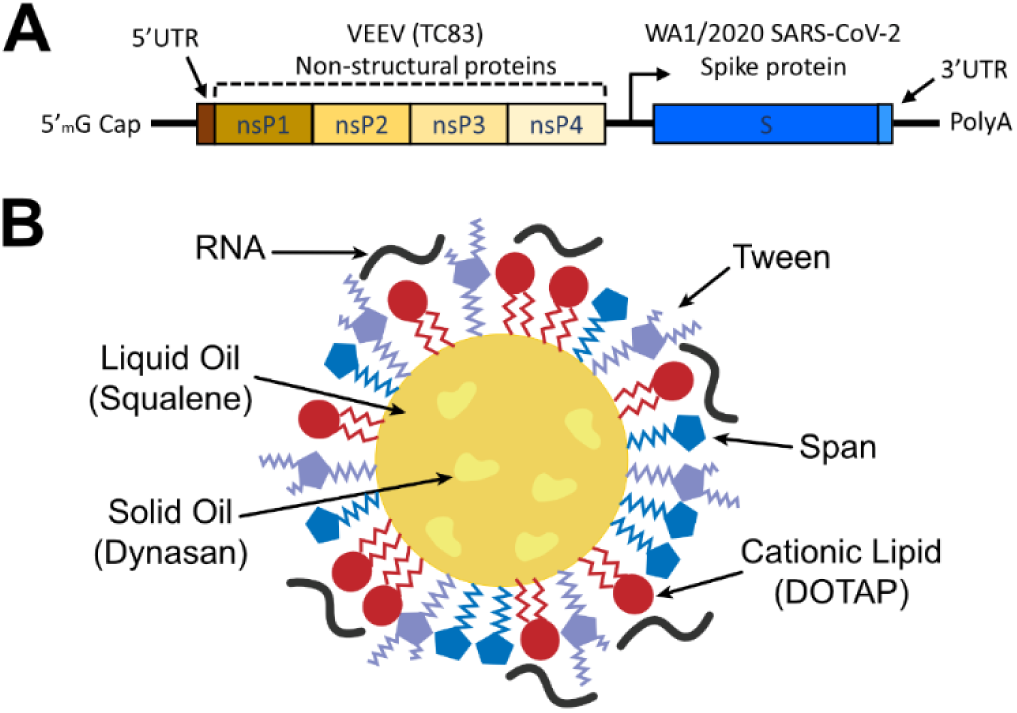
Design of saRNA-NLC SARS-CoV-2 vaccine AAHI-SC2. **(A)** saRNA SARS-CoV-2 construct diagram. **(B)** NLC-RNA complex diagram.

To investigate the ability of the saRNA-NLC vaccine to generate effective immunogenicity when delivered intranasally, 6–8-week-old C57BL/6J mice were given prime and boost vaccine doses of 1, 5, or 10 µg, 3 weeks apart. Control mice were i.n. dosed with 10 µg of a non-immunogenic saRNA-NLC expressing the reporter protein secreted alkaline phosphatase 2 (SEAP). Systemic immunogenicity was compared between groups. Serum, bone marrow, lung, and spleen tissues were harvested after vaccination and tested for humoral (serum IgG and pseudovirus neutralization titers; bone marrow antibody-secreting cells [ASCs] by enzyme-linked immunospot [ELISpot]) and cellular (systemic T cell responses by intracellular cytokine staining and flow cytometry [ICS-flow] and T cell ELISpot of splenocytes; mucosal T cell responses by ICS-flow of lungs) vaccine-induced SARS-CoV-2-directed immune responses.

The i.n. saRNA-NLC vaccine elicited high serum spike-specific IgG and SARS-CoV-2 Wuhan-strain neutralization titers that would be predicted to be protective (>1000 IC_50_) ^33,34^ at all i.n. doses after prime and boost (Figure 2A, B). Strikingly, at the 5 and 10 µg i.n. doses, no significant differences were observed from the 10 µg i.m. vaccines post-boost, suggesting equivalent induction of serum responses – the key correlates of protection for SARS-CoV-2 ^33,34^ – by the i.n. vaccine. IgG1 and IgG2a serum antibody titers indicated that i.n. vaccination with the saRNA-NLC vaccine maintains the desired Th1 bias seen with our i.m. vaccine (Figure 2C). Next, bone marrow-resident ASCs were assessed post-boost by ELISpot to determine if there was establishment of a memory niche for IgG-secreting ASCs. No differences were seen between the i.n. vaccines at either the 5- or 10-µg dose as compared to the 10 µg dose i.m. (Figure 2D), further suggesting effective systemic induction by both delivery routes and suggesting durability of response established by both vaccines.

**Fig. 2.**
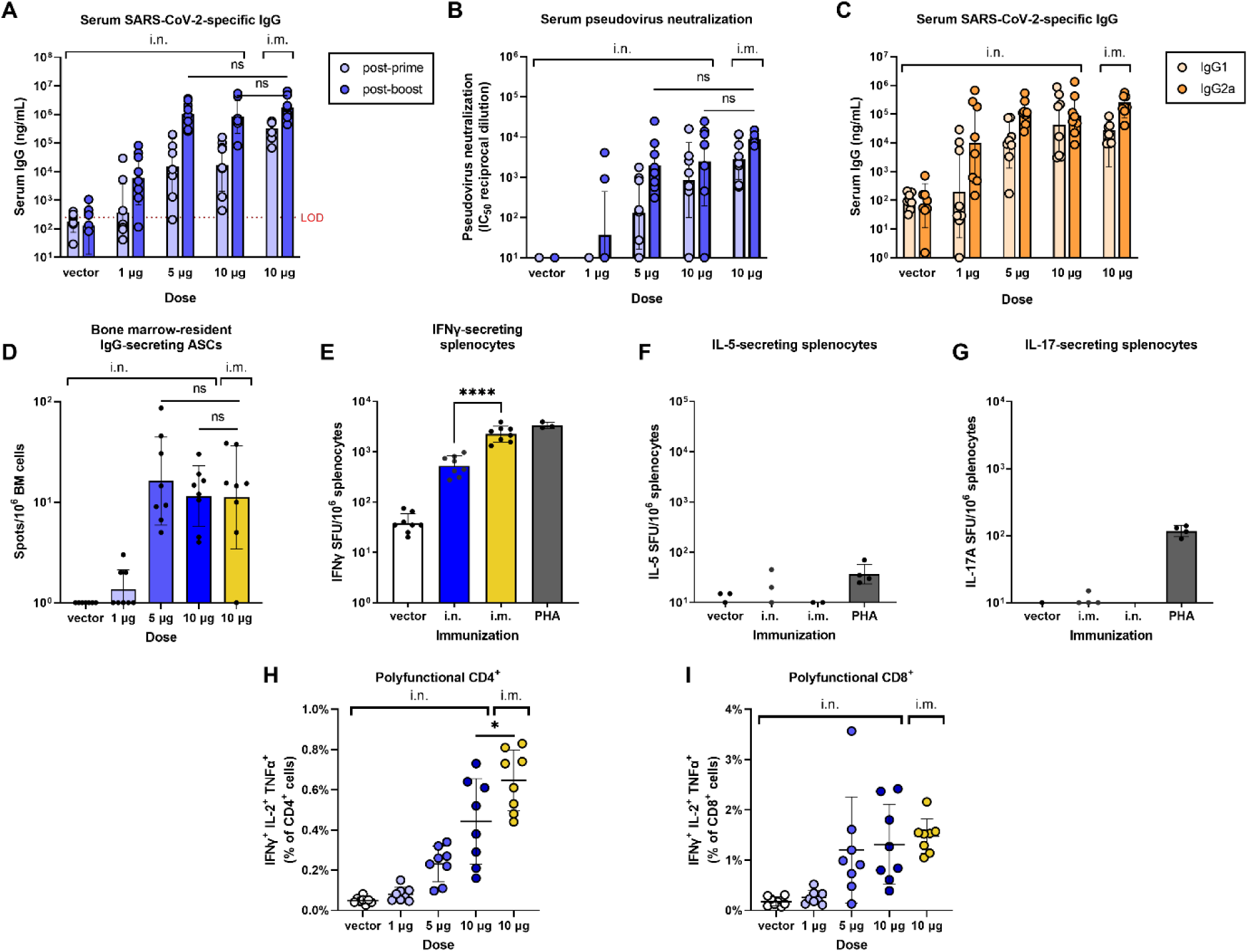
Systemic immunogenicity of intranasally administered SARS-CoV-2 saRNA-NLC vaccine. Systemic immunogenicity was assessed 3 weeks post-prime or 3 weeks post-boost in C57BL/6J mice immunized with AAHI-SC2 i.m. at 10 µg or i.n. at 1, 5, or 10 µg or a vector control (SEAP) i.n. at 10 µg at day 0 and day 21. *n* = 8 mice per group, evenly split male-female. **(A)** Serum SARS-CoV-2 spike-specific IgG responses measured post-prime and post-boost. LOD = limit of detection. **(B)** Serum SARS-CoV-2 spike Wuhan pseudovirus neutralizing titers measured post-prime and post-boost. **(A-B)** Data were log transformed and analyzed by a mixed-effects model with Tukey’s multiple comparisons test. **(C)** Serum spike-specific IgG1 and IgG2a responses measured post-boost. **(D)** Induction of bone marrow-resident spike-specific IgG-secreting ASCs as measured by ELISpot 3 weeks post-boost. **(E-G)** SARS-CoV-2 spike-responsive secretion of IFNγ, IL-5, or IL-17A by splenocytes post-boost as measured by ELISpot in mice dosed with 10 µg of saRNA. All bars on graphs show geometric mean ± geometric SD. **(H-I)** Splenic spike-responsive polyfunctional T cells (IFNγ^+^, IL-2^+^, and TNFα^+^ CD4^+^ or CD8^+^ T cells) post-boost as measured by intracellular cytokine staining and flow cytometry. Horizontal lines show mean ± SD. See Figure S1 for flow cytometry gating strategies. **(D-I)** Data were analyzed by one-way ANOVA with Tukey’s multiple comparisons test. \**p* < 0.05 and \*\*\*\**p* < 0.0001.

Cell-mediated immunity, particularly T cell immunity, is also thought to be critical to sustained, long-term protection against SARS-CoV-2 ^13,34^. We therefore investigated systemic cellular responses in i.n. and i.m. vaccinated mouse spleens after boost, both by ELISpot and ICS. Significant populations of IFNγ-secreting T cells were observed in both the i.n. and i.m. dosed (10 µg dose) mouse groups as measured by ELISpot (Figure 2E), indicating effective induction of effector T cells primed to respond to SARS-CoV-2. While the cell numbers were significantly higher in the i.m. dosed group, both groups showed robust T cell responses. IL-5- and IL-17A-secreting cells were seen at negligible populations in either group (Figure 2F-G), suggesting a strong Th1-biased response ^35,36^ induced by both the i.n. and i.m. vaccines with very little induction of a potentially harmful Th2 response associated with immunopathology and natural infection ^35,37,38^.

These robust, strongly Th1-skewed systemic CD4^+^ and CD8^+^ T cell responses were verified using ICS-flow performed on mouse splenocytes collected post-vaccination. After prime-boost immunization, significant populations of spike-reactive polyfunctional (IFNγ^+^, IL-2^+^, and TNFα^+^) CD8^+^ and CD4^+^ T cells were observed in mice immunized with the 5- and 10-μg i.n. vaccine doses and the 10-μg i.m. dose (Figure 2H, I), demonstrating strong T cell-mediated responses induced by both i.m. and i.n. vaccines. The spike-reactive polyfunctional T cell populations of CD4^+^ T cells induced by the i.n. vaccine were lower than those induced by the i.m. vaccine (20-30% reduction between the i.n. and i.m. vaccines at a 10-µg dose, *p* = 0.02); however, spike-reactive polyfunctional CD8^+^ T cell populations were not significantly different between the i.n. and i.m. vaccinated mice, and CD4^+^ T cell responses induced by the i.n. vaccine were still high, demonstrating effective seeding of systemic T cell immunity across all i.n. and i.m. vaccinated mice. As seen with vaccine-induced antibody titers, a stronger induction of a Th1-specific response (secretion of IFNγ^+^, IL-2^+^, or TNFα^+^) compared to a Th2 response (secretion of IL-5^+^ or IL-10^+^) or a Th17 response (secretion of IL-17A^+^) occurred within the T cell compartment in both i.n. and i.m. vaccinated mice and across all doses (Figure S2A), suggesting an overall systemic functional response to the virus. These data therefore indicate that the i.n. saRNA-NLC vaccine elicits humoral and cellular responses at levels predicted to be protective.

### Intranasal vaccine administration induces specialized lung T cell populations

While both i.m. and i.n. vaccines induce excellent systemic immunity, a key objective to assessing i.n. immunization was to determine whether effective mucosal responses could be generated with i.n. administration of an RNA vaccine. Immune responses in the lower respiratory tract are likely to be an important contributor to a vaccine’s ability to prevent severe disease ^39–41^. To that end, the lung T cell compartment was analyzed from C57BL/6J mice vaccinated i.n. or i.m. with 10 µg of AAHI-SC2 3 weeks post-prime and 3 weeks post-boost; the lung T cell compartment was analyzed by ICS-flow for the presence of specialized T cell subsets that would indicate the development of long-lasting respiratory-specific immunity.

First, we examined post-vaccination lung infiltration of antigen-experienced memory T cells. These T_RM_ cells are characterized by CD69 expression as well as CD103 expression in the CD8 compartment ^42^. The percentage of T_RM_ cells were significantly increased in the CD8 compartment with i.n. vaccination (median of CD69^+^ cells >20% of CD8^+^) compared to i.m. vaccination (median of CD69^+^ cells <10% of CD8^+^) following boost (CD69^+^ CD8^+^ cells, *p* = 0.003; CD69^+^ CD103^+^ CD8^+^ cells, *p* < 0.0001) (Figure 3A, B). While the fractions of CD8^+^ T_RM_ cells were equivalent post-prime, lung expansion of T_RM_ cells followed boost administration in the i.n. dosed group only, suggesting seeding of these durable populations followed by recall and expansion upon boost dosing.

**Fig. 3.**
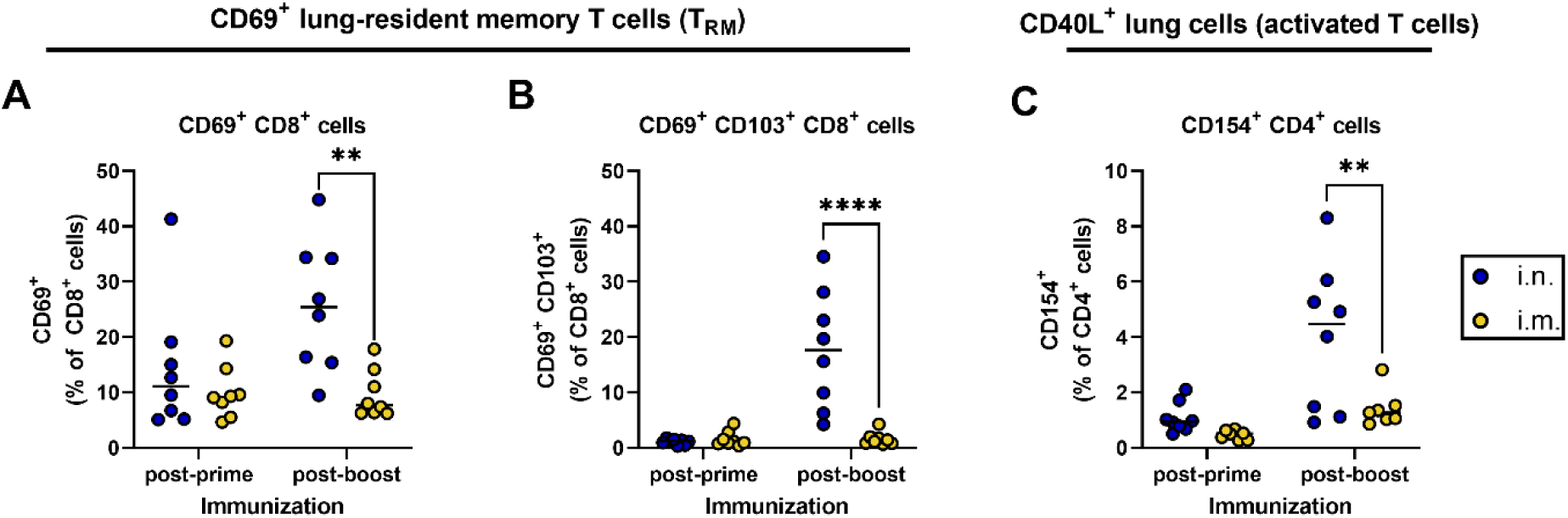
Mucosal T cell populations following intranasal vaccination with SARS-CoV-2 saRNA-NLC candidate. T cell populations in the lungs were assessed 3 weeks post-prime and 3 weeks post-boost in C57BL/6J mice immunized with 10 µg of AAHI-SC2 either i.m. or i.n. *n* = 8 mice per group, evenly split male-female. Vector control-dosed mice were also included in the experiment and are shown in Figure 4. See Figure S3 for sample flow cytometry gating strategies. **(A-B)** CD69^+^ lung-resident memory **(A)** CD8^+^ and **(B)** CD103^+^ CD8^+^ T cells. **(C)** CD40L^+^ (CD154^+^)-expressing CD4^+^ T cells. Percentage of cells in total live lung cells per mouse indicated by dots with horizontal line at median. Data were analyzed by two-way ANOVA with Sidak’s multiple comparisons test. \*\**p* < 0.01 and \*\*\*\**p* < 0.0001.

We then assessed CD154^+^ CD4^+^ T cells. CD154 (CD40 ligand [CD40L]) is often expressed on activated T cells and T follicular helper cells and is critical for licensing dendritic cells and engaging with B cells to enhance mucosal antibody production ^43^. Here, we found similar patterns to those seen with T_RM_ cells; significantly increased populations of CD154^+^ CD4^+^ cells were observed in the lungs after i.n. saRNA-NLC vaccination post-boost compared to i.m. vaccination (*p* = 0.003) (Figure 3C). Collectively, these data show that i.n. administration appears to elicit an expansion of T_RM_ and CD40L^+^ activated CD4^+^ T cells within the lung. In contrast, i.m. vaccination elicits minimal levels of these cell subsets, suggesting that these specialized respiratory cells are not inducible by i.m. vaccination.

### Intranasal administration elicits polyfunctional SARS-CoV-2 reactive mucosal T cell responses

While the lung T cell subsets were present in large populations post-i.n. administration, we wanted to verify that these cell subsets were functional effector cells and confirm that they responded to SARS-CoV-2. Here, we took mice that received the vector control or had been immunized and boosted with AAHI-SC2 i.m. (10 µg) or i.n. (1 µg, 5 µg, or 10 µg) and used ICS-flow cytometry to determine the frequency of polyfunctional lung IFNγ^+^IL-2^+^TNFα^+^ CD4^+^ and CD8^+^ T cells following re-stimulation for 6 hours with overlapping spike peptides.

Polyfunctional IFNγ^+^IL-2^+^TNFα^+^-secreting SARS-CoV-2 spike-responsive T cells composed a large fraction of the key T_RM_, CD69^+^ population in the 5- and 10-µg i.n. immunized mice (12.8% and 9.11%, respectively), which was significant in the CD69^+^ CD8^+^ subset of these T_RM_ cells compared to the i.m. immunized mice (*p* = 0.0005 for the 5-µg dose and *p* = 0.03 for the 10-µg dose), as well as in the CD69^+^ CD103^+^ CD8^+^ subset (*p* = 0.02) from 5-µg i.n. immunized mice (Figure 4A-B). Intramuscularly immunized groups again induced consistently negligible levels of polyfunctional CD8^+^ cells. We also examined the activated CD154^+^/CD40L^+^ CD4^+^ T cells and the lung-homing CD194^+^/CCR4^+^ CD4^+^ and CD8^+^ cells. Similar to the T_RM_ cells, CD154^+^ CD4^+^ cells showed significantly higher percentages of polyfunctional antigen-reactive cells at the 5 µg i.n. dose compared to the i.m. dose (*p* = 0.0006) (Figure 4C). Likewise, CD194^+^ CD4^+^ and CD8^+^ cells were also induced by the i.n. vaccine at significantly higher percentages at the 5 µg dose (*p* = 0.04 and *p* = 0.01, respectively) compared to the i.m. vaccine (Figure 4D-E).

**Fig. 4.**
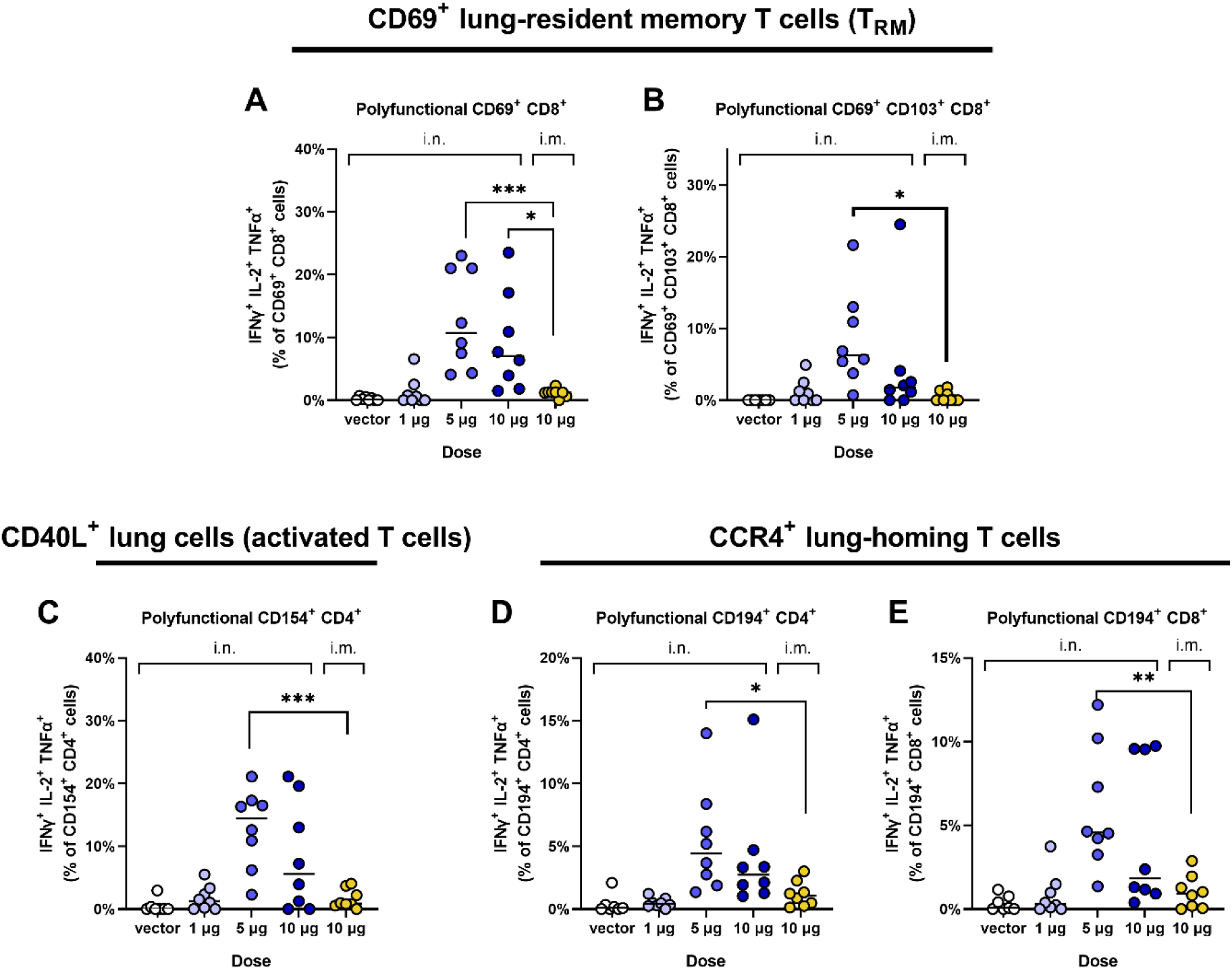
Mucosal SARS-CoV-2 spike polyfunctionally reactive T cell populations following immunization with the intranasal or intramuscular SARS-CoV-2 saRNA-NLC vaccine. T cell populations in lungs from C57BL/6J mice immunized with 1, 5, or 10 µg of the i.n. saRNA-NLC vaccine, 10 µg of the i.m. vaccine, or an i.n. vector control (SEAP) saRNA-NLC. T cells from day 21 post-prime were stimulated with SARS-CoV-2 spike peptides for 6 hours and stained for T cell markers and cytokine expression 3 weeks post-boost in. *n* = 8 mice for all dosing groups. Percentages of polyfunctional cells (expressing IFNγ, IL-2, and TNFα) were then measured by flow cytometry. **(A-B)** Polyfunctional CD69^+^ lung-resident memory (A) CD8^+^ and (B) CD103^+^ CD8^+^ T cells. **(C)** Polyfunctional CD40L-expressing (CD154^+^) CD4^+^ T cells. **(D-E)** Polyfunctional CCR4-expressing (CD194^+^) lung-homing **(D)** CD4^+^ and **(E)** CD8^+^ T cells. Dots represent values measured from individual mice. Horizontal lines indicate data median. Significance was determined by one-way ANOVA with Tukey’s multiple comparisons test: \**p* < 0.05, \*\**p* < 0.01, and *** *p* < 0.001.

Finally, we examined the quality of the response by lung T cells to confirm that i.n. administration elicited Th1-biased lung T cell responses, like what was observed in the spleen. The majority of the responses to both the i.n. and i.m. vaccines were Th1-specific responses, with negligible Th2 and Th17 responses (Figure S2B), similar to what was observed in the spleen. Th2 responses were not significantly different between i.m. vaccinated mice and mice receiving the vaccine via the i.n. route, did not scale with increasing vaccine dose, and were not significantly increased from buffer-only groups. Collectively, these data demonstrate that i.n. administration seeds specialized polyfunctional CD4^+^ and CD8^+^ T cell populations in the lung – populations which are not elicited following i.m. administration.

### Intranasal boosting of previous i.m. vaccination induces both systemic and mucosal immune responses

To understand whether i.n. administration could be viable as a boosting strategy for individuals previously vaccinated with an i.m. SARS-CoV-2 vaccine, we examined mice that were primed with an i.m. vaccine (10 µg) and then boosted with a medium dose (5 µg) either i.m. or i.n. Intranasally boosted mice showed equivalent serum anti-spike IgG levels and similar systemic neutralizing antibody responses to the i.m. boosted mice (Figure 5A, B). Within the splenic CD4^+^ and CD8^+^ T cell compartment, i.n. boost immunization induced equivalent or increased spleen-resident polyfunctional T cell responses in CD4^+^ or CD8^+^ (*p* = 0.05) cells, respectively, compared to an i.m. boost immunization (Figure 5C, D), indicating that an i.n. booster may effectively boost the systemic immunity developed in response to an original i.m. vaccination. With respect to mucosal immunity, the i.n. booster induced a higher, although not significant, percentage of CD40L^+^ (CD154^+^) activated CD4^+^ T cells within the lungs than the i.m. booster (Figure 5E). Most importantly, the i.n. booster elicited significantly more polyfunctional lung-resident CD8^+^ CD69^+^ T_RM_ cells (*p* = 0.02) than the i.m. booster (Figure 5F), indicating that simply boosting with an i.n. dose may be sufficient to elicit the full mucosal immune-stimulating effect of i.n. vaccination. These data are consistent with other i.n. vaccines that suggest i.n. boosting can have enhanced efficacy derived from mucosal targeted immunization as compared to i.m. vaccines ^21^. Lastly, to confirm that polyfunctional CD69^+^ T cells induced by i.n. prime+boost and i.m. prime + i.n. boost immunization were indeed lung-resident, we injected CD45 intravenously to discriminate vascular (CD45^+^) and alveolar (CD45^-^) T cells (Figure S4). As expected, i.n. immunization delivered during the prime+boost or only in the booster immunization resulted in higher frequencies of polyfunctional CD45^-^ CD69^+^ CD4^+^ and CD8^+^ T cells compared to i.m. prime+boost immunization alone (Figure S4A and D). Moreover, a significantly greater proportion of polyfunctional CD69^+^ CD4^+^ and CD8^+^ T cells were CD45^-^ following i.n. administration (Figure S4B-C and E-F). Taken together, these results suggest that an i.n. boost vaccination strategy may be ideal to boost systemic responses and induce respiratory immunity in previously vaccinated and/or infected individuals, reflecting the real-world scenario of second-generation SARS-CoV-2 vaccine rollouts.

**Fig. 5.**
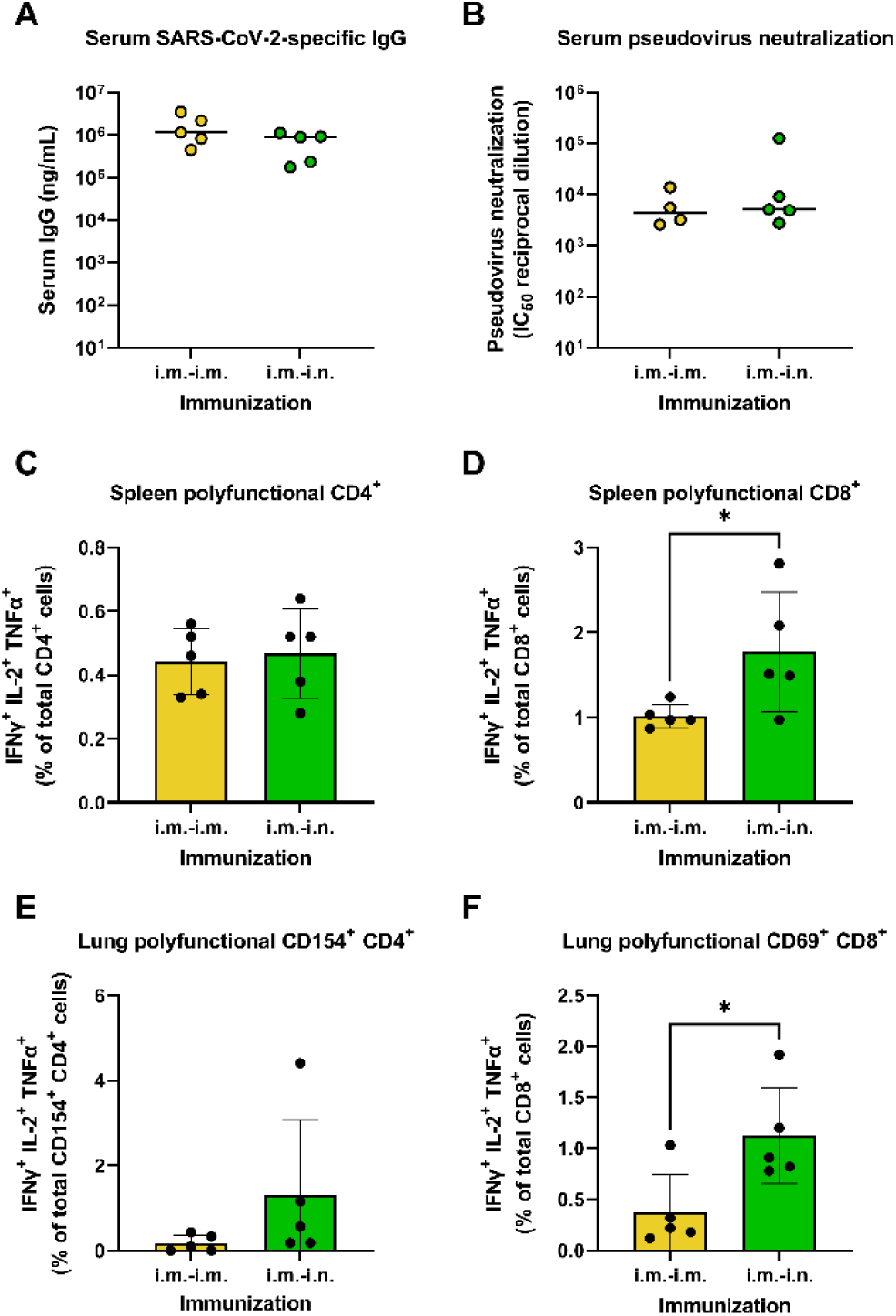
Systemic and mucosal immune responses following boost immunization with the intranasal or intramuscular SARS-CoV-2 saRNA-NLC vaccines. 10-week-old C57BL/6J mice were injected i.m. with a 10-µg prime dose and then boosted with a 5-µg dose of either the i.n. or i.m. saRNA-NLC vaccine 7 weeks later. *n* = female 5 mice per group. **(A)** SARS-CoV-2 spike-specific IgG titers as measured by ELISA 3 weeks post-boost. Horizontal lines show median. **(B)** Serum SARS-CoV-2 pseudovirus neutralizing titers 3 weeks post-boost. Horizontal lines show median. **(C-D)** Spleen polyfunctional **(C)** CD4^+^ and **(D)** CD8^+^ T cell responses. **(E)** Polyfunctional CD40L-expressing (CD154^+^) CD4^+^ T cells. **(F)** Polyfunctional lung-resident memory CD69^+^ CD8^+^ T cell responses. Statistics evaluated as unpaired two-sided homoscedastic *t* tests, \**p* < 0.05. Bars on graphs show mean with SD.

### saRNA-NLC vaccination protects against SARS-CoV-2-induced disease in hamsters

Given our data showing that i.m. prime and i.n. boost with our saRNA-NLC SARS-CoV-2 vaccine elicited strong systemic and pulmonary immunity in mice, we hypothesized that vaccination, particularly via the i.m./i.n. route, would be sufficient to protect animals from morbidity due to SARS-CoV-2 infection and would prevent these animals from transmitting the virus to naive bystander animals. To test this hypothesis, we co-housed hamster pairs (1:1) for several weeks to acclimate them to one another. Then, paired animals were separated and one animal from each pair was either sham-vaccinated or vaccinated with 3 µg of the saRNA-NLC SARS-CoV-2 vaccine given either i.m. or i.n (Figure 6A). After 24 hours, the sham-vaccinated and vaccinated animals were re-paired with their naive (non-vaccinated) cagemates. On day 21, peripheral blood was collected from all animals. On day 28, all pairs of animals were separated, and the sham-vaccinated and vaccinated animals were boosted via the i.m. or i.n. route. All animals were re-paired with their naive cagemates after 24 hours and blood was collected from all animals five days later. As expected, vaccinated hamsters, particularly i.m.-dosed hamsters, demonstrated strong induction of spike-specific serum IgG post-prime, which was absent in the sham-vaccinated group (Figure 6B) and in all the naive cagemates (Figure S5). Intranasally primed animals also showed development of significant serum IgG titers, albeit at titers lower than those observed in i.m.-primed animals (*p* < 0.0001). Boosting on day 28 did not result in a significant increase in systemic anti-spike IgG responses in any of the vaccinated groups (Figure 6C), perhaps due to the still ongoing robust primary response.

**Fig. 6.**
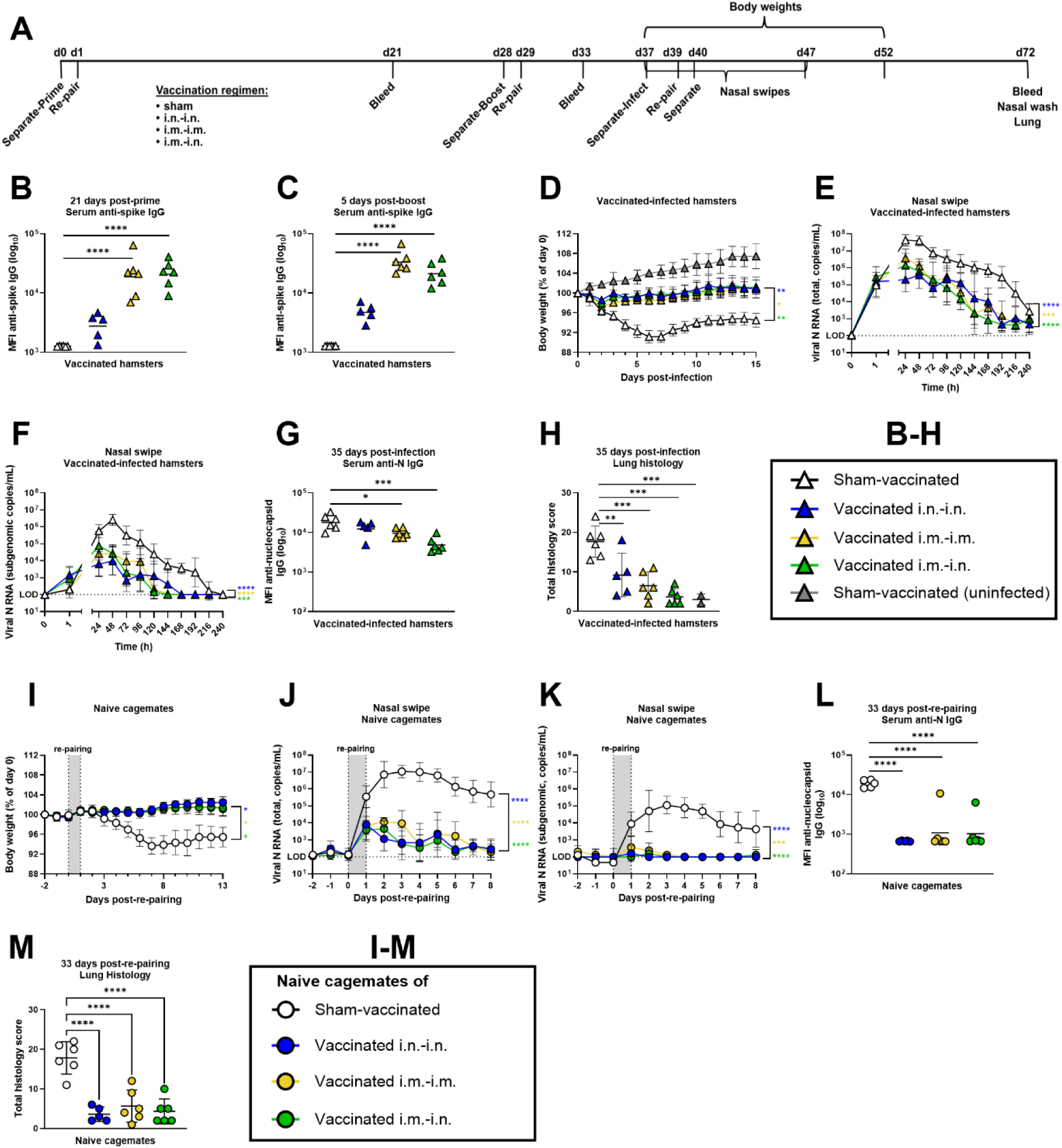
Vaccination of hamsters with the SARS-CoV-2 saRNA-NLC prevents SARS-CoV-2 infection-induced morbidity and prevents transmission of live virus to naïve hamster cagemates. **(A)** Study design to measure protection and viral transmission by SARS-CoV-2 saRNA-NLC prime+boost vaccinated hamsters that were challenged 9 days following the boost with live SARS-CoV-2. **(B-C)** Vaccine-elicited antibody responses in sham-vaccinated (open black triangles), i.n.-prime/i.n.-boost vaccinated (blue triangles), i.m.-prime/i.m.-boost (yellow triangles) or i.m.-prime/i.n.-boost vaccinated (green triangles) animals (*n* = 5-6/group). Serum anti-spike IgG levels reported as log-transformed values of the mean fluorescence intensity (MFI) of antibody binding to SARS-CoV-2 spike-coated beads on **(B)** day 21 post-prime and **(C)** 5 days post-boost (day 33). Significance assessed by two-way ANOVA with Tukey’s multiple comparison test. Horizontal lines show geometric mean. **(D-H)** Immune responses and viral load in sham-vaccinated and prime+boost vaccinated hamsters challenged 9 days following the boost with live SARS-CoV-2. **(D)** Body weight of prime+boost vaccinated-infected, sham-vaccinated-infected, and sham-vaccinated-uninfected hamsters. Data shown as group weight mean and SEM. Longitudinal data analyzed as area under the curve and assessed statistically using one-way ANOVA with Tukey’s multiple comparison test. **(E)** Total and **(F)** subgenomic nucleocapsid (N) RNA copy number in nasal swipes collected longitudinally over 10 days post-infection in prime+boost vaccinated-infected and sham-vaccinated-infected hamsters. The log-transformed viral RNA copy number for each group and timepoint is reported as geometric mean and geometric SD. Area under the curve measurements were assessed using one-way ANOVA with Tukey’s multiple comparison test. **(G)** Anti-N IgG levels in serum isolated on day 35 post-infection from prime+boost vaccinated-infected and sham-vaccinated-infected animals reported as log-transformed MFI values of binding to SARS-CoV-2 N protein-coated beads. Data analyzed using one-way ANOVA with Dunnett’s multiple comparisons test. Horizontal lines show geometric mean. **(H)** Lung pathology scores calculated for sham-vaccinated-infected and prime+boost vaccinated-infected animals 35 days post-infection. Horizontal lines and error bars represent mean and SD. **(I-M)** Immune responses and viral load in naïve hamsters following re-pairing with sham-vaccinated-infected (open black circles), i.n.-prime/i.n.-boost vaccinated (blue circles), i.m.- prime/i.m.-boost vaccinated (yellow circles), or i.m.-prime/i.n.-boost vaccinated (green circles) cagemates (*n* = 5-6/group). **(I)** Body weight of the naïve hamsters before and following re-pairing with prime+boost vaccinated-infected or sham-vaccinated-infected hamsters. Data shown as group weight mean and SEM. Longitudinal data analyzed as area under the curve using one-way ANOVA with Tukey’s multiple comparison test. **(J)** Total and **(K)** subgenomic viral N copy number measured in nasal swipes collected longitudinally over 8 days in naïve hamsters following re-pairing with prime+boost vaccinated-infected and sham-vaccinated-infected cagemates. The log-transformed viral RNA copy number for each group and timepoint is reported as geometric mean and geometric SD. Data assessed using one-way ANOVA with Tukey’s multiple comparison test. **(L)** Anti-N IgG levels in serum from naive hamsters isolated on day 33 post-re-pairing with prime+boost vaccinated-infected and sham-vaccinated-infected cagemates. Antibody levels reported as log-transformed MFI and analyzed using one-way ANOVA with Dunnett’s multiple comparisons test. Horizontal lines show geometric mean. **(M)** Lung pathology scores calculated in naive hamsters on 33 days post-re-pairing with sham-vaccinated-infected and prime+boost vaccinated-infected cagemates. Horizontal lines and error bars represent mean and SD, respectively. * *p* < 0.05, ** *p* < 0.01, *** *p* < 0.001, **** *p* < 0.0001. See also Figures S5-S7.

At 9 days post-boost (day 37 post-prime on timeline in Figure 6A), the vaccinated and sham-vaccinated hamsters were separated from their naive cagemates and experimentally infected with 1 x 10^6^ pfu SARS-CoV-2 (USA/WA-1 strain) delivered intranasally. Some sham-vaccinated hamsters were set aside to serve as uninfected negative controls (sham-vaccinated-uninfected). We then compared morbidity of the vaccinated-infected and sham-vaccinated-infected animals to the sham-vaccinated-uninfected controls. As expected, the sham-vaccinated-uninfected hamsters exhibited a modest increase in body weight over the next 14 days (Figure 6D). In contrast, the sham-vaccinated-infected animals had increased morbidity with peak body weight loss of 9-10% by day 6 post-infection and minimal body weight recovery by termination of study. The vaccinated-infected animals (i.n.-i.n., i.m.-i.n., and i.m.-i.m. regimens), on the other hand, showed minimal weight loss and exhibited rapid weight recovery by day 3-4 post-infection.

Importantly, the vaccinated-infected animals had significantly reduced virus infection-induced morbidity when compared to their sham-vaccinated-infected counterparts (*p* = 0.01 for i.n.-i.n., *p* = 0.01 for i.m.-i.n., and *p* = 0.02 for i.m.-i.m.). Body weight loss after challenge did not correlate with systemic anti-spike IgG titers (Figure S6), suggesting that all three vaccination regimens induced sufficient systemic IgG to protect against SARS-CoV-2-induced weight loss and/or that cellular or mucosal responses played a role in protection.

To further understand the dynamics of infection and transmission, total and subgenomic viral nucleocapsid (N) RNA was measured in nasal swipes from all infected hamsters across the first 10 days after infection. Total viral N RNA was detected in all infected hamsters as early as 1 hour post infection (Figure 6E), which confirmed successful intranasal instillation of SARS-CoV-2. Total viral copies peaked between 24-48 hours in all infected groups. However, total viral N RNA copies were significantly lower in the vaccinated-infected groups compared to the sham-vaccinated-infected group (*p* < 0.0001 for i.n.-i.n., *p* < 0.0001 for i.m.-i.n., and *p* = 0.0003 for i.m.-i.m.). Next, subgenomic viral N RNA was quantified in the nasal discharge to compare infectious, actively replicating viral load between the sham-vaccinated-infected and vaccinated-infected hamsters. Subgenomic viral N RNA was detected in the nasal swipes from all infected hamsters. However, significantly diminished subgenomic viral N RNA copies were observed in all vaccinated-infected animals relative to the sham-vaccinated-infected animals (Figure 6F) (*p* < 0.0001 for i.n.-i.n., *p* = 0.0002 for i.m.-i.n., and *p* < 0.0001 for i.m.-i.m.). Moreover, the peak copies of subgenomic N RNA in nasal swipes were observed within 24 hours of infection in the vaccinated-infected hamsters, while subgenomic N RNA peaked in the sham-vaccinated-infected hamsters between days 1-2 (Figure 6F). Importantly, the vaccinated-infected hamsters had increased clearance of viral subgenomic N RNA that declined to undetectable levels by day 6 (i.m.-i.n. and i.m.-i.m. groups) and day 8 (i.n.-i.n. group) post-infection. In contrast, the sham-vaccinated-infected control animals had detectable viral subgenomic N RNA until day 10, indicating that vaccination promoted more rapid reduction in replicating virus load in the respiratory tract.

The viral RNA measurements from nasal swipes suggested that none of the saRNA-NLC vaccination regimens induced sterilizing immunity following direct intranasal instillation with 1×10^6^ pfu of SARS-CoV-2. To address whether the vaccinated animals made an immune response to experimental infection, we measured systemic anti-N IgG antibodies on day 35 post-infection. Indeed, anti-N IgG antibody was detected in the serum from all three groups of vaccinated-infected animals (Figure 6G), indicating that the amount of N protein made by the infectious virus was sufficient to elicit a *de novo* N-specific antibody response in these hamsters. However, compared to the sham-vaccinated-infected animals, the amount of anti-N IgG antibody was significantly lower in the i.m.-i.m. and i.m.-i.n. vaccinated-infected hamsters (*p* < 0.0001 and *p* = 0.05, respectively) and modestly lower in the i.n.-i.n. vaccinated-infected hamsters, again consistent with a decreased viral load in all vaccinated groups of animals.

Next, to assess SARS-CoV-2-induced pulmonary pathology, cross sections of the hamster lungs were examined and scored for histology by a board-certified pathologist. As expected, the sham-vaccinated-infected hamsters had mild to moderate histology scores confirming that the SARS-CoV-2 viral inoculum used to infect the hamsters caused detectable pulmonary immunopathology as late as day 35 post-infection (Figure 6H). These sham-vaccinated-infected hamsters exhibited signs of inflammation that consisted primarily of accumulation of mononuclear and polymorphonuclear infiltrates in the alveolar interstitium and lumen, and adjacent inflammation around blood vessels and bronchioles (Figure S7A). Conversely, all three groups of vaccinated-infected hamsters had significantly decreased total histology scores (Figure 6H) and exhibited only sporadic interstitial, perivascular, and peribronchiolar inflammation (Figure S7B) that were comparable to sham-vaccinated-uninfected negative control animals (Figure S7C), suggesting that vaccination reduced the extent and severity of lung pathology following infection with a high dose of SARS-CoV-2. Therefore, while vaccination did not prevent infection following a high-dose viral challenge, all vaccine regimens effectively prevented disease-associated morbidity as measured by weight loss, decreased peak viral copies in the nasal discharge, suppression of viral replication, increased viral clearance, and protection against SARS-CoV-2-induced lung damage.

### saRNA-NLC vaccination prevents viral transmission by virally challenged hamsters

Although all three groups of vaccinated-infected hamsters appeared to have been productively infected following experimental SARS-CoV-2 exposure, the amount of subgenomic viral N RNA in the nasal secretions of these hamsters had already declined significantly by day 2 post-infection relative to the sham-vaccinated-infected group (Figure 6F). To assess whether this reduction in infectious viral load was sufficient to prevent transmission, we re-paired the sham-vaccinated-infected and vaccinated-infected hamsters with their naive cagemates on day 2 post-infection (day 39 post-prime on timeline in Figure 6A) for a period of 24 hours and assessed morbidity, nasal swipe viral RNA, serum IgG antibodies, and lung immunopathology in the naïve animals that were naturally infected through exposure to infected cagemates. As expected, naive cagemates of the sham-vaccinated-infected animals began losing weight within 24 hours of re-pairing, reaching maximum weight loss of 6-7% by day 7 post-re-pairing and exhibiting minimal body weight recovery by termination of study (Figure 6I). In striking contrast, the naive cagemates of the vaccinated-infected hamsters exhibited no body weight loss following re-pairing (*p* = 0.04 for i.n.-i.n., *p* = 0.05 for i.m.-i.n., and *p* = 0.05 for i.m.-i.m.).

Given the little morbidity observed in the naive cagemates co-housed with the vaccinated-infected hamsters, we hypothesized that viral transmission between the vaccinated-infected animals and their naive cagemates would also be diminished. To test this, we measured total (Figure 6J) and subgenomic (Figure 6K) viral N RNA in the nasal swipes from the naive cagemates that were paired for 24 hours with the sham-vaccinated-infected and vaccinated-infected groups. Total viral N RNA, which reflects both replicating and non-replicating virus, was present in nasal swipes from all the naive cagemates as early as 24 hours post-re-pairing with infected hamsters (Figure 6J). Interestingly, the naive cagemates paired with vaccinated-infected hamsters had on average 70-fold lower total viral N RNA at 24 hours post-re-pairing compared to naive hamsters that were co-housed with sham-vaccinated-infected animals (Figure 6J). Hence, immediately following re-pairing, the viral load transmitted to naive cagemates by the vaccinated-infected animals was lower than that transmitted by the sham-vaccinated-infected animals. Importantly, total viral N RNA in nasal secretions from the naive cagemates paired with vaccinated-infected hamsters never surpassed 10^4^ copies/mL (Figure 6J) and remained just above the limit of detection starting at day 4 post-re-pairing (*p* < 0.0001 for all three groups).

Conversely, the naive cagemates paired with sham-vaccinated-infected hamsters had total viral N RNA nasal discharge titers that peaked at 10^7^ copies/mL by day 2 post-re-pairing with minimal clearance by end of study at day 8 post-re-pairing (Figure 6J). Similarly, subgenomic viral N RNA in the nasal discharge, which is indicative of viral replication/infection, was detected at 24 hours post-re-pairing in the naive cagemates co-housed with sham-vaccinated-infected hamsters, peaked at 10^5^ copies/mL by day 2-3 post-re-pairing and remained detectable even out to day 8 post-re-pairing (Figure 6K). This was in striking contrast to the naive cagemates of the vaccinated-infected animals, which had minimal to non-detectable levels of subgenomic viral N RNA in the nasal discharge (Figure 6K) (*p* < 0.0001 for all three groups).

To assess whether the naive cagemates exposed to the vaccinated-infected animals were protected from active infection, we measured *de novo* anti-N serum IgG responses. As expected, the naive cagemates co-housed with sham-vaccinated-infected hamsters developed high serum N-specific IgG responses (Figure 6L). The naive cagemates co-housed with vaccinated-infected hamsters, on the other hand, developed negligible anti-N antibody responses (*p* < 0.0001 for all vaccine regimen groups). Indeed, no anti-N IgG response was detected in the naive cagemates of the i.n.-i.n. vaccinated-infected group, and only one naive cagemate from the i.m.-i.n and the i.m.-i.m. vaccinated-infected groups developed a measurable anti-N IgG serum antibody response. Thus, all three vaccine regimens were effective at preventing significant transmission of infectious virus capable of inducing an immune response to naive cagemates.

Next, we compared the total histology score of each group of naive cagemates. Naive hamsters co-housed with the sham-vaccinated-infected hamsters exhibited mild to moderate histology scores (Figure 6M), primarily due to mild inflammation in the alveolar interstitium and lumen that was similar in severity to that observed in the sham-vaccinated-infected hamsters (Figure S7A). Naive hamsters co-housed with the vaccinated-infected hamsters, regardless of vaccination regimen, exhibited a significantly decreased histology score (*p* < 0.0001 for all vaccine regimen groups), with minimal lung pathology that was very similar to that observed in the vaccinated-infected cagemates (Figure S7B). Taken together, the data demonstrate that i.m.-i.m., i.n.-i.n., or i.m.-i.n. dosing of the saRNA-NLC spike vaccine was not only effective in preventing disease-associated morbidity in hamsters but also prevented the vaccinated and subsequently experimentally infected hamsters from transmitting infectious virus to naive cagemates, even at a time when viral load peaked in the nasal discharge.

### saRNA-NLC vaccination induces long-term protective immunity and transmission prevention in hamsters

We next tested the durability of the protective immunity induced by saRNA-NLC vaccination. To do this, we selected the two prime-boost vaccine regimens that induced the most robust spike-specific IgG responses, namely i.m.-i.m. and i.m.-i.n. (Figures 6B and 6C), and measured protective immunity up to 3 months post-boost. Hamsters were either sham-vaccinated or vaccinated i.m. with 5-µg saRNA-NLC SARS-CoV-2 vaccine and then re-paired with naïve (non-vaccinated) cagemates (Figure 7A). On day 21 post-prime, nasal swipes and peripheral blood were collected from the vaccinated and sham-vaccinated animals. On day 28 post-prime, paired animals were separated again for 24 hours, during which time the sham-vaccinated and vaccinated animals were boosted with a 5 µg vaccine dose via the i.m. or i.n. route prior to re-pairing with the same naïve cagemates. Peripheral blood was collected again on day 55 post-boost, while nasal swipes were collected again between days 33 and 71 post-boost (Figure 7A). As expected, the sham-vaccinated animals did not develop anti-spike IgG or IgA antibodies (Figure 7B-F). By contrast, vaccinated animals seroconverted and exhibited increased levels of serum spike-specific IgG (Figure 7B) and IgA (Figure S8A) 21 days post-prime, which persisted for at least 55 days post-boost (Figure 7C and Figure S8B). spike-specific IgG antibodies were also detected in nasal secretions of vaccinated hamsters as early as day 21 post-prime (Figure S8C) and remained significantly elevated at 33 days post-boost (Figure S8D) and 71 days post-boost (Figure S8E) (*p* < 0.001 for both groups of vaccinated hamsters). No spike-specific IgA was detected in nasal swipes on day 21 post-prime in either sham-vaccinated or vaccinated animals (Figure 7D); however, increased levels of spike-specific IgA were detected in the nasal discharge at 33 days post-boost from both groups of vaccinated hamsters relative to sham-vaccinated controls (Figure 7E). Interestingly, spike-specific IgA levels in nasal swipes remained elevated up to 71 days post-boost in the i.m.-i.n. vaccinated animals but decreased to background control levels in the i.m.-i.m. vaccinated animals (Figure 7F), suggesting that an intranasal saRNA-NLC vaccine boost generates longer-lasting mucosal immunity than an intramuscular boost.

**Fig. 7.**
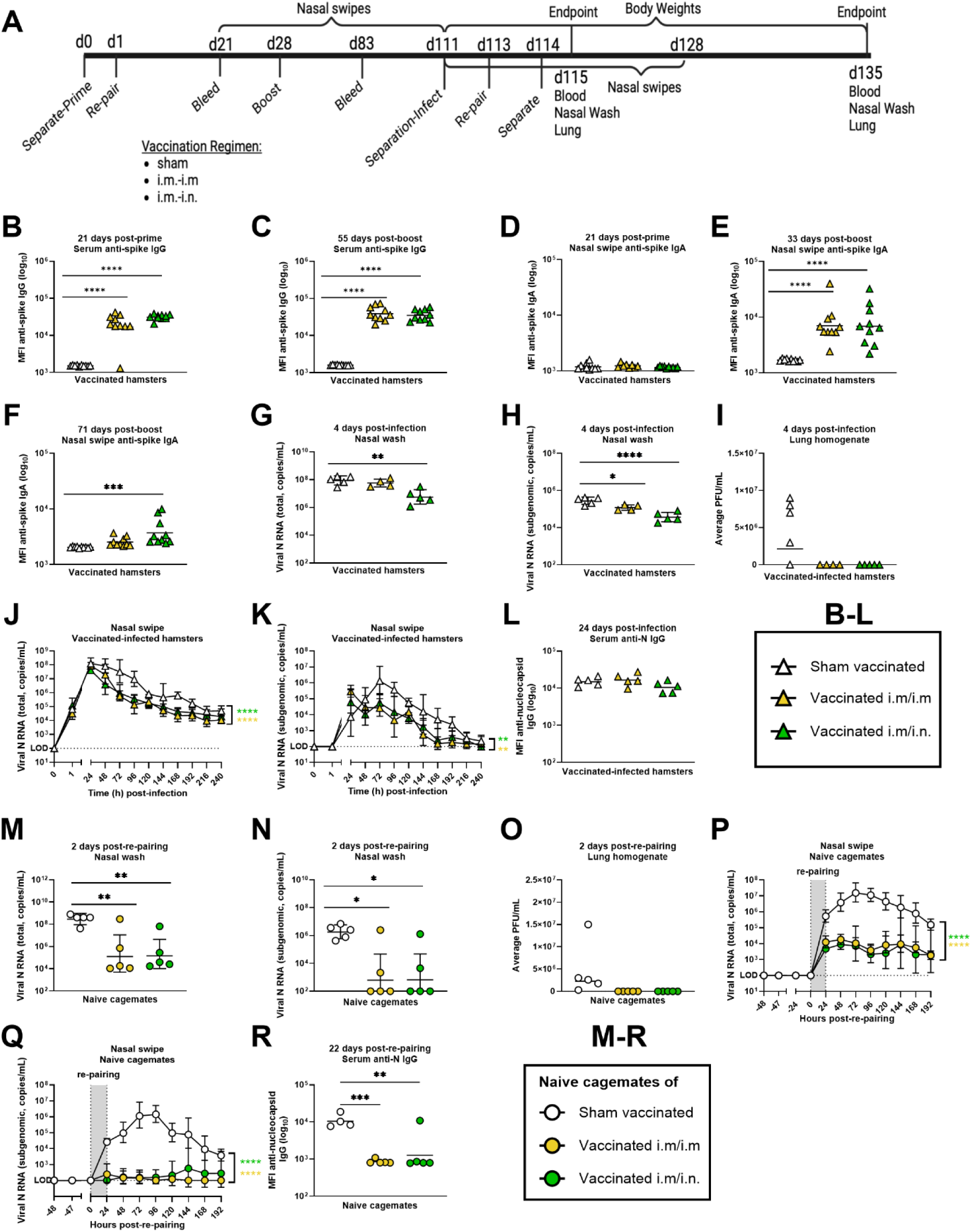
SARS-CoV-2 saRNA-NLC vaccination induces long-term protective immunity in hamsters and prevents viral transmission to naïve cagemates. **(A)** Study design to measure protection and viral transmission by SARS-CoV-2 saRNA-NLC prime+boost vaccinated hamsters that were challenged 83 days following the boost with live SARS-CoV-2. **(B-F)** Vaccine antibody responses in sham-vaccinated (open white triangles), i.m.-prime/i.m.-boost vaccinated (yellow triangles), or i.m.-prime/i.n.-boost vaccinated (green triangles) animals (*n* = 8-10/group). Anti-spike **(B-C)** IgG and **(D-F)** IgA levels reported as log-transformed MFI of binding to SARS-CoV-2 spike-coated beads. Serum anti-spike IgG levels reported on **(B)** day 21 post-prime and **(C)** 55 days post-boost (day 83). Nasal swipe anti-spike IgA levels reported on **(D)** day 21 post-prime, **(E)** 33 days post-boost (day 61), and **(F)** 71 days post-boost (day 99). Data analyzed using one-way ANOVA with Tukey’s multiple comparison test. Horizontal lines show geometric mean. **(G-L)** Immune responses and viral load in sham-vaccinated and prime+boost vaccinated hamsters infected 83 days post-boost (day 111) with SARS-CoV-2 (*n* = 4-5/group). **(G)** Total and **(H)** subgenomic N RNA copy number in nasal wash collected on day 4 post-infection of prime+boost vaccinated-infected and sham-vaccinated-infected hamsters. The log-transformed viral RNA copy number for each group is reported as geometric mean and geometric SD and analyzed using one-way ANOVA with Tukey’s multiple comparison test. **(I)** Infectious virus, measured by plaque assay, in lung homogenates on day 4 post-infection in prime+boost vaccinated-infected and sham-vaccinated-infected hamsters. Plaque forming units (PFU) reported as geometric mean of each group and analyzed using the Kruskal-Wallis test. **(J)** Total and **(K)** subgenomic N RNA copy number in nasal swipes collected longitudinally over 10 days in prime+boost vaccinated-infected and sham-vaccinated-infected hamsters. The log-transformed viral RNA copy number for each group and timepoint is reported as geometric mean and geometric SD. Area under the curve measurements assessed with one-way ANOVA with Tukey’s multiple comparison test. **(L)** Anti-N IgG levels in serum 24 days following infection of the prime+boost vaccinated-infected and sham-vaccinated-infected animals. Data reported as log-transformed MFI values of antibody binding to SARS-CoV-2 N protein-coated beads. Data analyzed using one-way ANOVA with Tukey’s multiple comparison test. Horizontal lines show geometric mean. **(M-R)** Immune responses and viral load in naïve hamsters following re-pairing with sham-vaccinated-infected (open white circles), i.m.-prime/i.m.-boost vaccinated (yellow circles), or i.m.-prime/i.n.-boost vaccinated (green circles) animals (*n* = 4-5/group). **(M)** Total and **(N)** subgenomic N RNA copy number measured in nasal wash from naïve hamsters 2 days following re-pairing with vaccinated-infected or sham-vaccinated-infected cagemates. The log-transformed viral RNA copy number for each group is reported as geometric mean and geometric SD and analyzed using one-way ANOVA with Tukey’s multiple comparison test. **(O)** Infectious virus, measured by plaque assay, in lung homogenates isolated from naive hamsters 2 days following re-pairing with vaccinated-infected and sham-vaccinated-infected cagemates. Plaque forming units (PFU) reported as geometric mean of each group and analyzed using the Kruskal-Wallis test. **(P)** Total and **(Q)** subgenomic N RNA copy number measured in nasal swipes collected longitudinally from naïve hamsters over 8 days following re-pairing with prime+boost vaccinated-infected or sham-vaccinated-infected hamsters. The log-transformed viral RNA copy number for each group and timepoint is reported as geometric mean and geometric SD. Area under the curve measures assessed with one-way ANOVA with Tukey’s multiple comparison test. **(R)** Anti-N IgG levels in serum from naïve hamsters 22 days following re-pairing with prime+boost vaccinated-infected or sham-vaccinated-infected cagemates. Data reported as log-transformed MFI values of binding to SARS-CoV-2 N protein-coated beads. Data analyzed using one-way ANOVA with Tukey’s multiple comparison test. Horizontal lines show geometric mean. * *p* < 0.05, ** *p* < 0.01, *** *p* < 0.001, **** *p* < 0.0001. See also Figure S8.

At 83 days post-boost (day 111 post-prime on timeline in Figure 7A), the vaccinated and sham-vaccinated hamsters were separated from their naïve cagemates and experimentally infected with 1 x 10^6^ pfu SARS-CoV-2 USA/WA-1 strain delivered intranasally. On day 4 post-challenge, one cohort of sham-vaccinated-infected and vaccinated-infected animals were euthanized for measurement of viral load and antigen-specific antibodies in the upper and lower respiratory tract. As expected, sham-vaccinated-infected animals had high total viral N RNA (Figure 7G) and subgenomic viral N RNA (Figure 7H) levels in the nasal wash, which is indicative of ongoing viral replication in the upper respiratory tract. Interestingly, total viral N RNA copies were significantly lower in i.m.-i.n. vaccinated-infected animals (*p* < 0.001) but not in the i.m.-i.m. vaccinated-infected counterparts compared to sham-vaccinated-infected controls (Figure 7G). However, both groups of vaccinated-infected animals had decreased subgenomic viral N RNA in the nasal wash compared to sham-vaccinated-infected animals (Figure 7H) (*p* < 0.05 for i.m.-i.m. and *p* < 0.0001 for i.m.-i.n.). Importantly, both groups of vaccinated-infected animals had detectable levels of spike-specific IgA in the nasal wash at 4 days post-infection (Figure S8F). We also measured viral load in whole lung tissue homogenates by plaque assay to directly quantify infectious virus particles in the lower respiratory tract. While geometric mean titer of 2.4 x 10^6^ pfu/mL was observed in lung tissue homogenates from sham-vaccinated-infected mice, no plaque-forming units were detected in the two groups of vaccinated-infected animals, confirming vaccine-induced protection from viral infection in the lung (Figure 7I).

A second cohort of experimentally infected hamsters were used for a longitudinal study to measure long-term vaccine efficacy against SARS-CoV-2 susceptibility and transmission. Hamsters challenged 83 days post-boost vaccination had decreased total (Figure 7J) and subgenomic (Figure 7K) viral N RNA copies in nasal swipes compared to sham-vaccinated controls, and exhibited moderate protection from SARS-CoV-2-induced body weight loss (Figure S8G). Similar levels of anti-N IgG antibodies were detected in the circulation of sham-vaccinated-infected and vaccinated-infected animals confirming a similar antibody response to the infectious SARS-CoV-2 virus in all three groups of experimentally challenged hamsters (Figure 7L).

Lastly, we sought to test whether the long-term protective immunity elicited by the saRNA-NLC SARS-CoV-2 vaccine would prevent viral transmission to naive cagemates. To do this, we re-paired the sham-vaccinated-infected and vaccinated-infected hamsters with their naive cagemates on day 2 post-challenge (day 113 on timeline in Figure 7A) for a period of 24 hours. By 2 days post-re-pairing, the naïve animals co-housed with the sham-vaccinated-infected animals were infected and presented with many copies of both total viral N RNA (10^8^-10^9^ copies/mL) (Figure 7M) and subgenomic viral N RNA (10^6^-10^7^ copies/mL) (Figure 7N) in the nasal wash. Importantly, the naïve animals co-housed with vaccinated-infected animals had decreased total (Figure 7M) and subgenomic (Figure 7N) viral N RNA compared to the animals paired with the sham-vaccinated-infected animals (*p* < 0.001 for total viral N RNA and *p* < 0.05 for subgenomic viral N RNA), suggesting that the vaccine significantly decreased viral transmission. To determine whether vaccine-induced protection extended to the lower respiratory tract, we measured viral titers 2 days post-re-pairing in lung tissue homogenates from the animals that were co-housed with the sham-vaccinated-infected animals and the vaccinated-infected animals. Animals co-housed with sham-vaccinated-infected animals had low but detectable infectious viral particles as measured by plaque assay, which were absent in the animals co-housed with vaccinated-infected animals (Figure 7O). No spike-specific IgA was detected in the nasal wash 2 days post-re-pairing in the animals paired with sham-vaccinated-infected or vaccinated-infected animals (Figure S8H), suggesting that 2 days following a natural SARS-CoV-2 infection is insufficient time to generate immunity in the nasal mucosa. Total and subgenomic viral N RNA was also measured daily in nasal swipes from animals co-housed with sham-vaccinated-infected and vaccinated-infected animals for 8 days post-re-pairing (Figures 7P and 7Q). Total viral N RNA was detected in animals co-housed with sham-vaccinated-infected animals as early as 24 hours post-re-pairing, peaking at 10^7^ copies/mL by 72 hours (3 days) and remaining above 10^5^ copies/mL at 192 hours (8 days) post-re-pairing (Figure 7P). Conversely, hamsters co-housed with vaccinated-infected animals had total viral N RNA copies that did not exceed 10^4^ copies/mL. The kinetics of subgenomic viral N RNA in nasal swipe of animals co-housed with sham-vaccinated-infected animals was similar to that of total viral N RNA, peaking above 10^6^ copies/mL by 3 days and still detectable at 8 days post-re-pairing (Figure 7Q).

Importantly, the animals co-housed with vaccinated-infected animals had low to undetectable subgenomic viral N RNA in nasal swipes (Figure 7Q) (*p* < 0.0001 for both groups of vaccinated-infected hamsters) indicating an almost complete absence of infectious virus in the nasal discharge. In addition, animals co-housed with vaccinated-infected animals had reduced body weight loss (Figure S8I) and negligible systemic N-specific IgG levels (Figure 7R) compared to animals co-housed with sham-vaccinated-infected animals, further highlighting the effectiveness of both the i.m.-i.n. and i.m.-i.m. vaccine regimens at inducing robust and long-lasting immunity that protects naive bystanders from viral transmission and disease-induced morbidity.

## DISCUSSION

Here we extend previous work on this saRNA-NLC vaccine platform to show delivery of an intranasal RNA vaccine that induces key respiratory mucosal immune responses. This i.n. administered saRNA-NLC vaccine induced systemic immunity in both the humoral and cellular compartments that rivaled i.m. administration. Additionally, i.n. administration elicited not only robust populations of spike-responsive polyfunctional T cells in the spleen but also in the lung, demonstrating induction of key respiratory mucosal T cell populations ^44–46^ by an RNA vaccine. Furthermore, we were able to demonstrate potential for this product as an i.n. booster of prior i.m. vaccine-induced immunity, a common real-life scenario given the largely COVID-experienced population more than 4 years from the start of the pandemic. This i.n. boost of prior i.m. vaccine-induced immunity significantly boosted systemic responses in addition to inducing mucosal T cell populations not stimulated by i.m. vaccine boosting. Lastly, we demonstrated that both i.n.-i.n., i.m.-i.m., and heterologous i.m.-i.n. vaccination regimens prevented virus-associated morbidity in the hamster model and, most importantly, prevented viral transmission and development of disease in naive cagemates when challenged either shortly after initial prime-boost vaccination, or even 3 months after boost vaccination.

With COVID-19 showing endemic trends post-peak pandemic and the number of patients suffering from long COVID continuing to rise ^47^, second-generation SARS-CoV-2 vaccines are urgently needed ^48^. While several vaccines have been developed and approved in the last 4 years, many utilizing novel mRNA vaccine technologies ^49–51^, which have admirably contributed to reduced disease mortality globally ^52^, emerging trends highlight the insufficiencies of current vaccine approaches. Vaccine efficacy has waned rapidly ^53–56^, and frequent boosters have been recommended for individuals that received Pfizer (Comirnaty) and/or Moderna (Spikevax) mRNA vaccines ^54,55,57^. Increasing evidence of reinfection, both post-infection and post-vaccination ^58^, particularly with Omicron and its sub-variants ^59^, as well as ongoing viral transmission post-vaccination ^60–62^, has negatively impacted vaccine credibility. With the threat of endemic SARS-CoV-2 circulation, the efficacy of these first-generation vaccines will likely not be sufficient to maintain control of existing and emerging viral strains ^63–65^.

While more research is necessary to understand why current vaccine responses wane quickly, emerging research points to insufficient nasal and mucosal immunity ^61,66^. Increasingly, data demonstrate that i.m. vaccination alone may not generate potent mucosal immunity that is predicted to be critical for early control and clearance of SARS-CoV-2 ^67^. Vaccinated individuals still show evidence of viral RNA in the upper respiratory tract after viral exposure, suggesting ongoing viral replication and transmission even in vaccinated populations ^20,22,61,68^. However, breakthrough infection with SARS-CoV-2 allows vaccinees to elicit potent mucosal and nasal immunity ^44–46^. Thus, improving the induction of nasal and mucosal immunity following vaccination is a critical component for next-generation vaccines to finally achieve durable and flexible protection. However, data also show that generating this immunity is generally not possible via standard i.m. vaccination and likely requires administration of an i.n. vaccine ^66,69^.

Intranasal vaccines to date have primarily been developed as live-attenuated flu vaccines (e.g., FluMist) ^70^. Recently, four new i.n. or inhalable vaccines have been approved for SARS-CoV-2 in China, India, Iran, and Russia, including two adenovirus-vectored vaccines ^71–73^. Further development of nasal or airway administration of vaccines for influenza and other respiratory viruses has remained an attractive target given the potential for development of critical airway T_RM_ cells at this first line of defense ^3,74^. Several other i.n. vaccines are in development for SARS-CoV-2, but most rely on standard protein or adenovirus vector vaccine technologies^20,21,46,75,76^. An RNA-based vaccine, in contrast, can be developed more rapidly, and allows cross-over i.m. to i.n. vaccination without developing a new vaccine, significantly increasing their utility in pandemic situations. The thermostability and stockpiling ability of our saRNA-NLC vaccine platform additionally provide key attributes for rapid and widespread pandemic response ^11,17^. Validating an RNA-based, i.n. administered vaccine technology would be a significant development not only for SARS-CoV-2 but for other respiratory pathogens, combining the rapidity and reliability of RNA vaccine adaptation to new targets with the key mucosal immune responses induced by mucosal-specific dosing.

AAHI’s SARS-CoV-2 saRNA-NLC vaccine candidate elicits both robust systemic humoral immunity and spike-responsive polyfunctional T cells as an i.m. or i.n. vaccine. Delivered i.n., it also elicits tissue-resident T cell populations within the lungs, suggesting broad T cell presence across the respiratory tract – an effect that i.m. delivered mRNA vaccines have been unable to induce to date ^44–46^. While CD4^+^ and CD8^+^ T cells are now being recognized as a major determinant of disease progression and critical to long-lasting immunity ^13^, the main vaccines on the market elicit minimal peripheral T cell responses ^77,78^ and negligible mucosal/respiratory T cell responses ^44–46^. The thermostability and ability to administer i.n. may make this vaccine substantially more desirable in the fight against respiratory viruses. These data demonstrate the potential for this saRNA-NLC vaccine candidate as an effective i.n. booster to pre-existing SARS-CoV-2 immunity. Indeed, unlike the i.m. booster, i.n. boosts elicit local durable mucosal immunity, suggesting that the additive effect of this i.n. vaccine to pre-existing systemic immune responses generated the greatest benefit of all vaccination regimens tested.

These data demonstrate that all immunization methods elicited full protection against virus-associated morbidity and viral transmission. While our original hypothesis was that only i.n. vaccination would prevent transmission, we were pleasantly surprised that all three delivery methods provided prevention of transmission, even when the challenge occurs months after the last dose for the i.m.-i.m. and i.m.-i.n. dosed vaccines tested in a durability study. These data bolster the overall strength of this saRNA-NLC vaccine platform. While it is unclear which specific cellular or humoral components mediated this protection in i.m. vaccinated animals, which lacked substantial lung T_RM_ cell populations and had reduced mucosal IgA compared to animals receiving one i.n. vaccination, we suspect that sufficiently high systemic IgG so early after boost (9 days post-boost) may continue to circulate and prevent transmission ^79,80^.

Protection and transmission were also examined in vaccinated hamsters at longer timepoints, along with infection and housing with naïve cagemates, which demonstrated the durability of both protection and the reduction of transmission by both i.m.-i.m. or heterologous i.m.-prime, i.n.-boost vaccine regimens. This may indicate that a sufficiently strong systemic vaccination can provide responses sufficient to protect against transmission even in the absence of mucosal immunity, though future studies will further investigate other mechanisms that may contribute to this protection.

The i.n. saRNA-NLC vaccine may be most effective as a booster dose following an i.m. prime. This conclusion aligns with the results of other studies ^14,68,81^, where priming an immune response with a strong i.m. vaccine followed by i.n. vaccination is able to optimally stimulate the T_RM_ compartment while still boosting the extant i.m. induced immunity. A body of studies have demonstrated the value of “prime-pull” vaccination strategies, where a strong “primed” systemic response can be “pulled” to a local, protective environment after an appropriately localized boost vaccination ^82,83^. Our data demonstrate the clear potential of using an identical or similar RNA vaccine product for both i.m. and i.n. vaccination, potentially eliminating many manufacturing hurdles and using an already clinically tested vaccine platform technology for multiple routes of administration ^32^. In our studies, i.n. boosting outperformed i.m. boosting in mucosal immune measures, hinting at the potential for i.n. dosing to access novel cell compartments and mechanisms. Further work remains to fully optimize the i.n. saRNA-NLC vaccine formulation specifically for i.n. administration, potentially including use of nasal spray devices, with the goal of enhancing induction of immunity while avoiding toxicity and reducing heterogeneity in nasal delivery. As a first step in this further development process, this i.n. SARS-CoV-2 vaccine candidate is being tested in nonhuman primate studies to enable future human testing.

These studies were carefully designed and carried out to maximize scientific rigor and reproducibility, however the studies do have certain limitations. We showed robust systemic and mucosal immunity, protection, and transmission prevention; these initial studies conducted in mice and hamsters may not translate fully to primates and humans. Ongoing studies have been designed to investigate the immunology and efficacy of this vaccine in non-human primates. Our data are also limited by the heterogeneity in mouse immune responses when vaccinated i.n., possibly related to a lack of consistent dosing devices for i.n. vaccination. Similarly, because mice are obligate nasal breathers ^84^ and vaccine volumes are difficult to appropriately scale to mouse size, some potential exists for delivery of a fraction of the vaccine volume beyond the nasal tract to the lungs and digestive tract. These issues are likely only resolvable with larger animal models better representing the nasal passages and respiratory tracts of humans, and use of appropriate delivery devices for improvement in vaccine dose delivery. Similarly, some toxicity was noted in the highest i.n. delivered saRNA-NLC vaccine dose (10 µg) in mice, resulting in transient mouse weight loss. While these effects are not expected to extend to primates due to non-linear dose scaling of saRNA vaccines relative to animal size and nasal tissue surface area, any potential toxicity noted with an i.n. vaccine will be closely monitored in future studies, and further optimization of the NLC formulation specifically for i.n. delivery will also be conducted to minimize any such effects. In the long-term hamster study, we note that changes in body weight may additionally be affected to age and stress associated with infighting as part of extended co-housing^85,86^, driving potential confounding of this readout. Finally, we were unable to test this saRNA-NLC vaccine candidate head-to-head against any commercially available SARS-CoV-2 vaccines for comparison of either vaccine immunogenicity or efficacy at the time of study. These comparator groups will support future investigations, including enabling further exploration of the use of this i.n. saRNA-NLC vaccine as an effective mucosal immunity-inducing booster of prior approved SARS-CoV-2 vaccination.

This saRNA-NLC vaccine platform represents a promising technology to combat both the existing SARS-CoV-2 pandemic and other respiratory illnesses, such as seasonal and pandemic influenza. For respiratory pathogens, i.n. vaccination or i.n. vaccine boosting of i.m. vaccination may help raise RNA vaccine technologies to a new level, through stimulating respiratory T cell and B cell populations ^27,87,88^ and generating effector subsets within the lungs and nasal mucosa primed to respond rapidly at the first point of infection ^68^. Beyond its efficacy at stimulating T_RM_ subsets and bolstering systemic immunity, i.n. administration may also be desirable as a needle-less delivery method with potentially higher uptake, particularly in pediatric and vaccine-hesitant populations. Altogether, this novel demonstration of a unique i.n. delivered RNA vaccine that elicits strong systemic and mucosal immunity highlights a promising new pandemic response technology ripe for future development and application.

## METHODS

### Study design

#### Research objectives

This study was designed to test the immunogenicity and efficacy of i.n. and i.m. delivered SARS-CoV-2 saRNA-NLC vaccines in mice and hamsters. Our hypothesis prior to the study initiation was that i.n. delivery of this saRNA vaccine would elicit superior respiratory immunity in rodents, specifically eliciting long-lived memory T cells, while possibly driving systemic immune responses, even if not at the level induced by standard i.m. delivery of the vaccine.

Furthermore, we believed that the i.n. delivered vaccine would therefore elicit protection from morbidity in the hamster model and superior reduction in viral transmission relative to the i.m. delivered vaccine. As the data developed, we further hypothesized that heterologous dosing of the vaccine, with an i.m. prime and i.n. boost, would combine the best attributes of both vaccine delivery methods and elicit a combination of systemic and respiratory immune responses that would optimally protect against viral challenge and transmission.

#### Research subjects or units of investigation

Research was conducted on C57BL/6J mice obtained from The Jackson Laboratory. Lakeview Golden (LVG) Syrian hamsters were obtained from Charles River Laboratories.

#### Experimental design

We conducted several mouse vaccine immunogenicity experiments to address these questions, vaccinating mice i.m and/or i.n. with our SARS-CoV-2 saRNA-NLC vaccine. Mice were vaccinated using different doses and dosing strategies (homologous vs. heterologous prime-boost vaccine regimens), and immunogenicity was assessed by ELISA and pseudoneutralization assay on serum samples, T cell intracellular cytokine staining (ICS) on spleen and lung samples, bone marrow B cell ELISpot, and T cell ELISpot to assess systemic antibody responses and induction of specific vaccine-induced T cell and B cell populations.

#### Randomization

Rodents were obtained commercially and randomized into study groups prior to study onset.

#### Blinding

Investigators were not blinded to the study groups during data collection and analysis as this is typically unnecessary for this type of inbred animal study and introduces more systemic and human error than it prevents. Quantitative, unbiased assays were developed with highly regulated SOPs in order to minimize data biases.

#### Sample size

Statistical analyses have been the guiding strategy to maximize statistical power while reducing the number of animals used to a minimum. We have previously determined that 6-8 animals per group (3-4 male, 3-4 female) is the lowest estimate to achieve meaningful comparisons between vaccine candidates with subtle differences in immunogenicity. 10 mice/group, 5 male and 5 female, is required to provide statistical power of 90% with an alpha (*p*-value) of 0.05 to detect a statistically significant difference in pseudovirus neutralizing antibody titers between vaccinated groups with two-fold differences in mean antibody titers with expected statistical variance. We use both male and female mice in most study groups to avoid sex-driven data biases and allow for identification of any sex-specific differences in vaccine immunogenicity. For one select study where mouse numbers were limited (Figure 5), *n* = 5 all-female mice were used per group to provide statistical power of 80% to detect two-fold differences in serum antibody titers. The use of all female mice in this study was conducted in order to reduce data variability sufficiently to minimize animal use without compromising the main study immunogenicity readouts.

#### Data inclusion/exclusion criteria

No data were excluded from any plots or analyses except two cases: (1) for flow cytometry after ICS where <50,000 cells were collected in the well (this was established as a guideline prospectively) and (2) one i.n. vaccinated hamster was removed from Figure 6 where the complete lack of anti-spike IgG antibody on day 21 post-prime and 5 days following the boost established that it had not been effectively vaccinated.

#### Outliers

Our policy prior to study initiation was not to exclude any outliers. Therefore, we have not excluded any outliers. Assays were repeated on any suspected outliers in order to verify data prior to inclusion.

#### Replicates

Replication of data was conducted whenever possible. All experiments were conducted with a minimum of biological triplicates. ELISAs and pseudoneutralization assays were conducted with technical duplicates for each study sample. T cell immunogenicity was only assessed in technical singlicate due to limiting tissue quantity and cell number. Bridging groups were used between independent animal studies to provide replication of key study groups across multiple studies offset by months. All attempts at replication verified the robustness of the scientific approach.

### saRNA expression plasmid design, cloning, and production

The saRNA plasmid was designed and selected based on immunogenicity measures from i.m. immunization ^11^. Briefly, the saRNA encodes a SARS-CoV-2 spike sequence based on a GenBank sequence (MT246667.1) containing the D614G mutation, a diproline at sites 987-988, and a QQAQ substitution for RRAR in the furin cleavage site (683-686). The sequence was codon-optimized for expression in humans, synthesized by BioXp, and inserted via Gibson cloning into AAHI’s backbone saRNA expression vector. A SEAP-expressing plasmid was also used as a control, created similarly with the SEAP sequence in the place of the vaccine antigen. Sanger sequencing was used to confirm plasmid sequences. These were then amplified in *Escherichia coli* and extracted using Qiagen maxi- or gigaprep kits, linearized with NotI (New England Biolabs), and purified using a Qiagen DNA purification kit.

### RNA manufacture

NotI-linearized DNA plasmids containing T7 promoters were used as templates for *in vitro* transcription (IVT) of saRNA for vaccination. Our in-house optimized IVT protocol with T7 polymerase, RNase inhibitor, and pyrophosphatase (Aldevron) was used. Next, a DNase step was used to digest the templates, and Cap0 structures were added to the RNA transcripts with guanylyltransferase (Aldevron), GTP, and S-adenosylmethionine (New England Biolabs).

CaptoCore 700 resin (GE Healthcare) was used to chromatographically purify the RNA followed by diafiltration and concentration. saRNA material was filtered through a 0.22 µm polyethersulfone filter and stored at −80°C until further use. saRNA size and integrity was characterized by gel electrophoresis, and RNA concentration was quantified by UV absorbance (NanoDrop 1000) and RiboGreen assay (Thermo Fisher).

### NLC manufacture

A mixture of trimyristin (IOI Oleochemical), squalene (Sigma-Aldrich), sorbitan monostearate (Sigma-Aldrich), and the cationic lipid 1,2-dioleoyl-3-trimethylammonium-propane (DOTAP; Corden) was heated at 70°C in a bath sonicator. Separately, polysorbate 80 (Fisher Scientific) was diluted in 10 mM sodium citrate trihydrate and heated to 70°C in a bath sonicator. After all components were in suspension, a high-speed laboratory emulsifier (Silverson Machines) running at 7,000 rpm was used to blend the oil and aqueous phases. The colloid mixture was then processed at 30,000 psi for 11 discrete microfluidization passes using an M-110P microfluidizer (Microfluidics). The NLC product was then filtered through a 0.22 µm polyethersulfone filter and stored at 2°C–8°C until use ^11,17,89^.

### Vaccine complexing and characterization

Vaccine complexes were produced by combining aqueous diluted RNA at a 1:1 volume ratio with NLC that was pre-diluted in a 10 mM sodium citrate and 20% w/v sucrose buffer. Vaccines were prepared at a Nitrogen:Phosphate (N:P) ratio of 5-15, representing the ratio of amine groups on the NLC DOTAP to the RNA’s backbone phosphate groups. This complexing resulted in vaccine containing the intended dose of complexed saRNA-NLC in an isotonic 10% w/v sucrose, 5 mM sodium citrate solution. The vaccine solution was incubated on ice for 30 minutes after mixing to ensure complete complexing.

### Mouse studies

All mouse work was done according to the Bloodworks Northwest Research Institute’s Institutional Animal Care and Use Committee. All mouse work was in compliance with all applicable sections of the Final Rules of the Animal Welfare Act regulations (9 CFR Parts 1, 2, and 3) and the *Guide for the Care and Use of Laboratory Animals* ^90^.

C57BL/6J mice purchased from The Jackson Laboratory were used for all mouse studies. Mice were between 6 and 8 weeks of age at study onset and evenly split between male and female. At arrival, mice were simultaneously randomized into study groups, half male and half female, housed separately 3-5 mice per cage with irradiated bedding in Allentown individually ventilated cages. Sterile water gel packs were given to all newly arrived animals during the acclimation period, and animals had access to fresh potable water *ad libitum* via Edstrom automatic reverse-osmosis and chlorinated water system. Cages were changed every 2 weeks inside a laminar flow cage change station. A spot check for dirty cages was conducted on the non-change weeks.

Mice were observed daily for 3 days post-injection after both prime and boost injections by trained personnel. Isoflurane delivered by inhalation was used for temporary anesthesia immediately prior to retro-orbital blood collection and i.n. immunization to mitigate animal pain and distress. At study completion, animals were humanely euthanized by CO_2_ overdose.

Animal handling technicians were not blinded to study groups. After 1 week of acclimatization to the facility, mice were then primed with vaccine. Mice were then boosted 3 weeks later and harvested 3 weeks after boost. Mouse weights were monitored for 4 days following prime and boost vaccinations. Mice were immunized by i.m. injection in both rear quadriceps muscles (50 µL/leg, 100 µL total) or by i.n. inoculation in two 25 µL doses. Intranasal inoculation was done under full anesthesia, and mice were allowed to recover under a heat lamp. During ongoing studies, serum samples were taken by retro-orbital bleed. Three-weeks post-boost, mice were harvested, terminal bleeds were taken, spleens and lungs were dissected, and femurs were harvested for bone marrow purification. For intravenous labeling of lung-resident T cells, mice were anesthetized with isoflurane delivered by inhalation, injected with 100 µL of a 1:100 dilution of BUV395 CD45 (BD Biosciences #565967) via retro-orbital injection, allowed to recover for 3 minutes for antibody circulation, and then euthanized for tissue harvest.

### Mouse serum IgG, IgG1, and IgG2a titers by ELISA

SARS-CoV-2 spike-specific IgG was measured by ELISA. Plates (Corning 384-well high-binding #CLS3700) were coated with Recombinant SARS-Cov-2 Spike His Protein, Carrier Free (R&D Systems #10549-CV), at 1 µg/mL and incubated overnight at 4°C. Plates were incubated in blocking buffer (2% dry milk, 0.05% Tween 20, 1% goat serum) for over 1 hour. Samples were plated on low-binding plates, starting at a dilution of 1:40 and then serially diluting 1:2 across the plate. High-binding plates were washed, and samples were then transferred from the low-binding plates to the coated, high-binding plates. Naive mice serum at 1:40 was used as a negative control. A SARS-Cov-2 neutralizing monoclonal antibody (GenScript #A02057) was used as a positive control at a starting concentration of 3.2 ng/µL. Following a 1-hour incubation, plates were washed, and a secondary antibody, Anti-Mouse IgG (Fc Specific)-Alkaline Phosphatase antibody (Sigma-Aldrich #A2429), at a 1:4000 dilution was added. Following the secondary antibody and another wash step, phosphatase substrate tablets (Sigma-Aldrich #S0942) were dissolved in diethanolamine substrate buffer (Fisher Scientific #PI34064) at a final concentration of 1 mg/mL and added to the plates. Plates were incubated for 30 minutes and then read spectrophotometrically at 405 nm. A 4-point logistic (4PL) curve was used to fit the antibody standard titration curve, and sample concentrations were interpolated off the linear region of the curve.

For IgG1 and IgG2a isotype-specific ELISAs, plates were coated and blocked as described above. For IgG1 and IgG2a, the standard curve was run using SARS-CoV-2 neutralizing antibodies (GenScript #A02055 and #BS-M0220, respectively). Sample dilution and incubation were identical to the total IgG curve, and plates were probed with IgG1- and IgG2a-specific secondary alkaline phosphatase (AP)-conjugated detection antibodies (Sigma-Aldrich #SAB3701172 and #SAB3701179, respectively) prior to development, reading, and quantification as described above.

### Pseudovirus neutralization assay

SARS-CoV-2 pseudovirus neutralizing antibody titers in mouse sera were measured via a pseudoneutralization assay ^91^. Lentiviral SARS-CoV-2 spike protein pseudotyped particles were prepared by co-transfecting HEK-293 cells (American Type Culture Collection CRL #11268) with plasmids containing a lentiviral backbone-expressing luciferase and ZsGreen (BEI Resources #NR-52516), lentiviral helper genes (BEI Resources #NR-52517, NR-52518, and NR-52519), or a delta19 cytoplasmic tail-truncated SARS-CoV-2 spike protein (Wuhan strain plasmids from Jesse Bloom of Fred Hutchinson Cancer Center). Cells were incubated for 72 hours at standard cell culture conditions (37°C, 5% CO_2_), and cell culture media were harvested and filtered through a 0.2-μm filter. Pseudovirus-containing supernatant was frozen until titering and use.

To perform the assay, plates were seeded with Human Angiotensin-Converting Enzyme 2 (hACE2)-expressing HEK-293 cells (BEI Resources #NR52511) and incubated overnight. Serum samples were pre-diluted 1:10 in media (Gibco DMEM + GlutaMAX + 10% fetal bovine serum [FBS]) and then serially diluted 1:2 for 11 total dilutions. These were then incubated with polybrene (Sigma-Aldrich #TR-1003-G) and pseudovirus for 1 hour. Then, the serum samples were added onto the hACE2 cells in duplicate and incubated at 37°C and 5% CO_2_ for 72 hours. To determine 50% inhibitory concentration (IC_50_) values, plates were scanned on a fluorescent imager (Molecular Devices ImageXpress Pico Automated Cell Imaging System) for ZsGreen expression. Total integrated intensity per well was used to calculate the percent of pseudovirus inhibition per well. Neutralization data for each sample were fit with a 4-parameter sigmoidal curve, which was used to interpolate IC_50_ values.

### Spleen, lung, and bone marrow cell harvest

Spleens were prepared in 4 mL of RPMI medium by manual maceration through a filter using the back of a syringe. Dissociated splenocyte samples were centrifuged briefly at 400 x g to pellet fat cells, and the supernatants containing lymphocytes were either transferred to 5 mL mesh-cap tubes to strain out any remining tissue debris or lysed with ammonium-chloride-potassium (ACK) lysing buffer. Cells were counted on a Guava easyCyte cytometer (Luminex) and plated in round-bottom 96-well plates at 1-2 x 10^6^ cells per well in RPMI + 10% FBS containing CD28 costimulatory antibody (BD Biosciences #553294) and brefeldin A. One of three stimulation treatments was added to each well: 0.0475% dimethyl sulphoxide [DMSO], 2 μg/μL spike peptide pool (JPT peptides #PM-WCPV-S-1), or phorbol myristate acetate (PMA)-ionomycin. Plates were incubated for 6 hours at 37 C with 5% CO_2._

Lung cells were isolated via enzymatic digestion using a gentleMACS Dissociator (Miltenyi Biotec). Lungs were dissociated in 4 mL of Hanks’ Balanced Salt Solution (HBSS) supplemented with 10% Liberase (MilliporeSigma), 10% aminoguanidine, 0.1% KN-62, and 1.25% Dnase. Lungs and enzymatic mix were added to a gentleMACS M tube (Miltneyi Biotec), and the m_lung_01.02 program was run. Samples were then incubated at 37°C and 5% CO_2_ for 30 minutes. Directly after, samples were run again on the gentleMACS Dissociator using the m_lung_02.01 program. The resulting slurry was added to 10 mL of RPMI medium and centrifuged for 5 minutes. The supernatant was discarded, and the cells were either filtered through a 5 mL snap-top tube or washed again in RPMI medium prior to counting on a Guava easyCyte cytometer and plated in round-bottom 96-well plates at 1-2 x 10^6^ cells per well in RPMI + 10% FBS, 50 μM beta-mercaptoethanol, CD28 costimulatory antibody, brefeldin A, and one of three stimulation treatments: 0.0475% DMSO, 2 μg/μL spike peptide pool (JPT peptides #PM-WCPV-S-1), or PMA-ionomycin. Plates were incubated for 6 hours at 37 C with 5% CO_2._

### Intracellular cytokine staining and flow cytometry

Following incubation, plates were centrifuged at 400 x g for 3 minutes, the supernatants were removed, and cells were resuspended in phosphate-buffered saline (PBS). Splenocytes were stained for viability with Zombie Green (Biolegend) in 50 μL of PBS. Cells were washed twice, and then spleen samples were incubated with CD16/CD32 antibody (Invitrogen #14-0161-86) to block Fc receptors. Next, the cells were surface stained in an additional 50 μL of staining buffer (PBS with 0.5% bovine serum albumin and 0.1% sodium azide). Spleen samples were stained with fluorochrome-labeled monoclonal antibody (mAb) specific for mouse CD4 (eBioscience #45-0042-82), CD8 (BD Biosciences #563068), CD44 (BD Biosciences #560568), and CD107a (BioLegend #121614), while lung samples were stained for CD4 (BioLegend #100526), CD8 (BD Biosciences #563068), CD44 (BD Biosciences #562464), CD69 (BioLegend #104508), CD103 (BioLegend #121435), CD194 (BioLegend #131220), and CD154 (BD Biosciences #745242). Cells were washed twice, permeabilized using the Fixation/Permeabilization Kit (BD Biosciences #554714) and stained intracellularly. Spleens were stained for TNFα (BioLegend #506327), IL-2 (BioLegend #503824), IFNγ (Invitrogen #25731182), IL-5 (eBioscience #12-7052-82), IL-10 (BD Biosciences #564081), and IL-17A (BD Biosciences #560820), while lungs were stained for TNFα (BioLegend #506327), IL-2 (BioLegend #503824), IFNγ (Invitrogen #25731182), IL-5 (BioLegend #504306), and IL-17A (BD Biosciences #560820). After two washes in staining buffer, cells were resuspended in 100 μL of staining buffer and analyzed using an LSRFortessa flow cytometer (BD Biosciences). After initial gating for live CD4^+^ or CD8^+^ lymphocytes, cells gating positive for all three activation markers IL-2, IFNγ, and TNFα were counted as activated polyfunctional T cells.

### Bone marrow harvest and ASC ELISpot

Antibody-secreting bone marrow-resident cell counts were measured by ELISpot. MultiScreen_HTS_ IP Filter plates (0.45 µm, MilliporeSigma) were treated with 15 µL of 35% ethanol for 30 seconds. Recombinant SARS-CoV-2 Spike His Protein, Carrier Free (R&D Systems #10549-CV-100), was diluted at 2 µg/mL in ELISpot coating buffer (eBioscience), and plates were coated with 100 µL. Plates were incubated overnight at 4°C. The next day, plates were washed with PBS with 0.1% Tween 20 and blocked with RPMI + 10% FBS for 2 hours.

Femurs were removed from mice and inserted into a snipped-end 0.6-mL Eppendorf tube and then inserted into a 1.5 mL Eppendorf tube with 1 mL of RPMI + 10% FBS. Femurs were centrifuged for 15 seconds at 10,000 rpm, and supernatant was discarded. Cell pellets were vortexed, resuspended in 200 µL of RBC Lysis Buffer (Invitrogen), and then incubated on ice for 30 seconds. An additional 800 μL of RPMI medium was added, and cells were centrifuged for 5 minutes at 400 x g before the supernatant was decanted. Cells were resuspended in 1 mL of RPMI + 10% FBS, counted, and transferred to prepared filter plates at 1 million cells per well followed by a 3-fold dilution across five adjacent wells.

Plates were incubated for 3 hours, then washed three times with PBS plus 0.1% Tween 20. Secondary antibody (Goat Anti-Mouse IgG-HRP or IgA-HRP [SouthernBiotech #1030-05 and #1040-05, respectively]) was added at a 1:1000 dilution in PBS with 0.1% Tween and 5% FBS overnight at 4°C. Plates were then washed three times in PBS with 0.1% Tween 20 and two times in PBS. 100 µL of Vector NovaRED Substrate Peroxidase (Vector Laboratories #SK-4800) was applied for 7 minutes to develop the plates. The reaction was quenched by rinsing plates with distilled water for 2 minutes, and plates were dried in the dark. Spots were counted and data were analyzed using ImmunoSpot v7 software (Cellular Technology Limited).

### T cell ELISpot assay

IFNγ (BD Biosciences #51-2525KZ), IL-17A (Invitrogen #88-7371-88), or IL-5 (BD Biosciences #51-1805KZ) at a 1:200 dilution in Dulbecco’s PBS (DPBS; Gibco) was used to coat ELISpot plates (MilliporeSigma). Plates were incubated overnight at 4°C and then washed and blocked with RPMI + 10% FBS for at least 2 hours. Previously harvested splenocytes (see above) were added at 2 x 10^5^ cells per well. Samples were then stimulated with 1 µg/mL PepMix SARS-CoV-2 (JPT Peptide Technologies #PM-WCPV-S-1). Plates were incubated at 37°C and 5% CO_2_ for 48 hours. Then plates were washed with PBS with 0.1% Tween 20. Next, detection antibody (IFNγ, BD Biosciences #51-1818KA; IL-17A, Invitrogen #88-7371-88; and IL-5, BD Biosciences #51-1806KZ) was diluted in ELISpot diluent (eBiosciences) at 1:250 and added overnight at 4°C. Plates were washed and then developed using Vector NovaRED Substrate Peroxidase for 15 minutes. The reaction was quenched with deionized water. Plates were then dried in the dark. Spots were counted and data were analyzed using ImmunoSpot.

### Hamster studies

Nine-week-old male Lakeview Golden (LVG) Syrian hamsters were purchased from Charles River Laboratories and maintained in the University of Alabama at Birmingham (UAB) animal facilities, which include the UAB Southeastern Biosafety Lab (SEBLAB) ABSL3 facilities. All hamster procedures were approved by the UAB Institutional Animal Care and Use Committee (IACUC protocol 22628) and the UAB Biosafety Committee (Protocol 22-160 and 22-161). All procedures were performed in accordance with National Resource Council guidelines.

### Hamster vaccination and infection

Adult male hamsters were housed in pairs and separated at the time of vaccination. One hamster from each pair was vaccinated with 3 (initial study) or 5 µg (durability study) of saRNA-NLC delivered i.m. (50 µL per rear quadriceps muscle, 100 µL total) or i.n. (25 µL/nare, 50 µL total). Sham-vaccinated hamsters received dPBS i.m. Vaccinated and sham-vaccinated animals were re-paired with naive cagemates after 24 hours. Cagemates were separated on day 28, and vaccinated animals were boosted with the same vaccine given via the i.m. or i.n. route. Animals were re-paired after 24 hours. Nine or 83 days after the boost, animals were separated, and the vaccinated and sham-vaccinated hamsters were infected i.n. with 1 x 10^6^ pfu SARS-CoV-2 USA/WA-1 variant (50 µL/nare, 100 µL total). Uninfected controls received 100 µL (50 µL/nare) of dPBS (vehicle). The infected animals were re-paired with the uninfected, unvaccinated naive cagemates at 48 hours post-infection. Cagemates were separated 24 hours later and remained apart until the end of the study. Hamster weights were measured before infection and daily until day 13-15 post-infection.

### Hamster blood collection and processing

Blood samples from sedated hamsters were collected from the lateral saphenous vein on day 21 post-prime and either 5 days or 55 days post-boost vaccinations. Blood samples following euthanasia were collected from terminal bleed via the posterior vena cava. Blood samples were transferred into BD Microtainer blood collection tubes (BD Biosciences), and serum was separated by centrifugation at 10,000 rpm at RT for 10 minutes. Serum samples collected post-infection were heated at 60°C for 20 minutes to inactivate SARS-CoV-2 virus. All samples were aliquoted and frozen at -80°C until analyzed by cytometric bead array (CBA).

### Hamster nasal swipes for viral RNA and antibody quantitation

Awake hamsters were transferred into an open container to minimize movement, and the exterior surface of their nose was swiped for 5-10 seconds with a polyester-tipped swab (Medical Packaging #SP-7D) that had been dipped in a screw-cap tube containing 100 µL (for CBAs) or 300 µL (for viral N RNA quantitation) viral transport medium (VTM; HBSS (+Ca^2+^ +Mg^2+^) containing 2% FBS, 100 mg/mL gentamicin, and 0.5 mg/mL amphotericin B). The tip of the swab was cut and placed inside the VTM-containing tube. The tubes were vortexed, and swab tips were discarded. Samples were then processed for viral RNA quantitation or virus-specific antibody CBAs.

### Hamster nasal wash samples

Hamsters were sedated with isoflurane and euthanized with an intraperitoneal injection of tribromoethanol (500 mg/kg, 3 mL/animal). The trachea was exposed, an incision was made, and a blunt 19-gauge needle attached to a 1-mL insulin syringe was inserted into the trachea. While holding the mouth of the hamster shut with one hand, 400 µL of VTM was squeezed through the nasal passages, out of the nose, and into a screw-cap tube. Samples were heated at 60°C for 20 minutes to inactivate SARS-CoV-2 virus, then aliquoted, and stored at -80°C until analyzed by CBA.

### Recombinant SARS-CoV-2 spike, RBD, and N protein production

Recombinant SARS-CoV-2 spike ectodomain trimeric proteins were prepared as previously described ^92^. Briefly, two human codon-optimized constructs were generated with a human IgG leader sequence, the SARS-CoV-2 spike ectodomain (amino acids 14-1211), a GGSG linker, T4 fibritin foldon sequence, a GS linker, and a 15 amino acid biotinylation consensus site (AviTag, construct 1) or 6X-HisTag (construct 2). Each construct was engineered with two sets of mutations to stabilize the protein in a pre-fusion conformation ^92^. Recombinant SARS-CoV-2 spike RBD domain monomers (human codon-optimized) were generated with a human IgG leader sequence, the spike RBD (amino acids 319-541 from SARS-CoV-2 Wuhan-Hu-1 strain), a GS linker, an AviTag, and a 6X-HisTag. The coding sequence of N from SARS-CoV-2 Wuhan-Hu-1 strain was synthesized in frame with the coding sequence for an AviTag and was cloned in frame to the 6X-HisTag in the pTrcHis2C expression vector (Invitrogen). Biotinylated recombinant N protein was produced by co-transforming Rosetta cells with the N expression plasmid and an inducible BirA expression plasmid. Bacteria were grown in the presence of chloramphenicol, ampicillin, and streptomycin, induced with IPTG, and supplemented with biotin. Recombinant SARS-CoV-2 Avi/His-tagged spike trimeric proteins were produced by co-transfecting each plasmid construct (1AVI:2HIS construct ratio) into FreeStyle 293-F cells. The Avi-His-tagged spike RBD protein was produced by transfecting the single dual-tagged RBD construct into FreeStyle 293-F cells. All recombinant proteins were purified by FPLC using a nickel-affinity column and subsequent size exclusion chromatography. After buffer exchange, purified spike ectodomain trimers and RBD domain protein were biotinylated by the addition of biotin-protein ligase (Avidity). Biotinylated proteins were buffer exchanged into PBS, sterile filtered, aliquoted, and stored at -80°C until used.

### SARS-CoV-2 spike cytometric bead array (CBA)

CBAs with recombinant SARS-CoV-2 proteins were prepared as previously described ^92^. Briefly, recombinant proteins were passively absorbed onto streptavidin functionalized 4 µm fluorescent microparticles (Carboxy Blue Particle Array Kit, Spherotech). 500 µg of biotinylated recombinant protein was incubated with 2 x 10^7^ streptavidin functionalized fluorescent microparticles in 400 µL of 1% BSA in PBS. Protein coupled beads were washed and resuspended at 1 x 10^8^ beads/mL and stored at 4°C.

### Cytometric bead array (CBA) measurement of spike-specific IgG and IgA responses

50 µL serum (diluted to 1/4000 in PBS), nasal wash, or nasal swipe (diluted 1/4 in PBS) was arrayed in 96-well u-bottom polystyrene plates. 5 µL of a suspension containing 5 x 10^5^ spike beads, RBD beads, and N beads was added to the samples. Suspensions were mixed, incubated for 15 minutes at RT, washed in PBS, stained with FITC-conjugated polyclonal anti-hamster IgG or PE-conjugated polyclonal anti-hamster IgA (Southern Biotech) for 15 minutes at RT, washed in PBS, and then resuspended in 100 µL of 1% paraformaldehyde in PBS. Samples were collected on a BD CytoFLEX flow cytometer in plate mode at a sample rate of 100 µL/minute for 1 minute. Following acquisition, the FCS files were analyzed in FlowJo (BD Biosciences). Briefly, the beads were identified by gating on singlet 4 µm particles in log scale in the forward scatter and side scatter parameters. APC-Cy7 channel fluorescence gates were used to segregate the particles by bead identity. Geometric mean fluorescent intensity was calculated in the FITC or PE channel.

### Propagation and titer determination of SARS-CoV-2

The original SARS-CoV-2 isolate USA-WA1/2020 was obtained from BEI Resources (#NR-52281), propagated in Vero E6 cells (ATCC #C1008), and titered as previously described ^93^. The handling of viable virus was performed in the Southeastern Biosafety Lab (SEBLAB) BSL-3 facility located at UAB.

### Viral RNA quantitation by qRT-PCR

SARS-CoV-2 viral N RNA and viral subgenomic N RNA was measured by PCR in nasal swipe (300 µL) samples as previously described ^93^. AccuPlex SARS-CoV-2 Reference Material (SeraCare Life Sciences, Inc., # 0505-0126) was extracted and amplified in parallel to generate a standard curve enabling viral N quantitation. The subgenomic RNA standard was constructed and validated in-house ^93^.

### Plaque assay

Six-well plates were seeded with Vero E6 cells (4E + 05/well) and incubated for 16 hours at 37°C and 10% CO_2_, and then serially diluted lung homogenate supernatants (1/10 in 1x PBS containing 1% FBS) were added to the cultures and incubated for 1 hour at 37°C with shaking every 10 minutes to allow for infection of cells by any infectious viral particles present in the samples. The virus solution was then aspirated, and the cells were overlayed with 0.6% Avicel solution supplemented with 3% FBS and 1x MEM. The plates were incubated in a CO_2_ incubator for 72 hours. The Avicel overlay was aspirated, and the cell monolayer was fixed with 10% neutral-buffered formalin for 1 hour. The cells were stained with 1% crystal violet solution and plaques were counted for each dilution. The virus titers were determined by multiplying the number of plaques by the dilution factor. The lowest limit of quantification for this assay was determined by the dilution of the lung tissue plated.

### Histopathology

Hamster lungs were inflated with 10% neutral-buffered formalin (Thermo Fisher) using a blunt 19-gauge needle. Each lung was excised *en bloc* and fixed for 7 days at RT in 15-fold volume of 10% neutral-buffered formalin. The fixed tissues were then embedded dorsal side down in paraffin using standard procedures. Samples were sectioned at 5 µm, and resulting slides were stained with hematoxylin and eosin (H&E). All tissue slides were evaluated by light microscopy by a board-certified veterinary pathologist blinded to study group allocations. Representative photo images were collected using a Nikon Eclipse Ci microscope (Nikon Inc) and analyzed with NIS-Elements software (Nikon Inc). Scoring was performed following a published algorithm ^94,95^. Briefly, the airway, alveolar, and vascular pathology in affected areas were assessed and scored on a scale of 0-4 ^93^, and then added together to generate a total histopathological score of the whole lung.

### Statistical analysis

The effect of dosing strategy on CD4 and CD8 cell polyfunctionality was assessed by one-way ANOVA with Tukey’s multiple comparisons test. The effect of dosing strategy on IgG titer and pseudovirus neutralizing antibody titers was assessed in log-transformed data using a mixed-effects model with Tukey’s multiple comparisons test. The effect of immunization strategy on IgG titer was assessed on log-transformed data using unpaired *t* tests or two-way ANOVA with Tukey’s or Sidak’s (if any sample data were missing) multiple comparisons test. The effect of immunization strategy on pseudovirus neutralizing antibody titers was assessed on log-transformed data using unpaired *t* tests or a mixed-effects model with Sidak’s multiple comparisons test. The effect of immunization strategy on polyfunctional CD4 and CD8 cell responses was assessed in untransformed data using unpaired *t* tests or one-way ANOVA with Holm-Sidak’s multiple comparisons test. Lung, bone marrow, and splenocyte ELISpot data were analyzed using log-transformed data by one-way ANOVA with Tukey’s multiple comparisons test. Hamster body weight data was analyzed using area under the curve measures assessed statistically by one-way ANOVA with Tukey’s multiple comparison test. Hamster viral load was analyzed using log-transformed area under the curve measures, assessed with one-way ANOVA with Tukey’s multiple comparison test. Plaque assays were reported as geometric mean of each group and analyzed using the Kruskal-Wallis test. All statistical analyses were conducted using Prism version 9 (GraphPad Software).

## Acknowledgments

We thank Jesse Bloom (Fred Hutchinson Cancer Center) for sharing the SARS-CoV-2 spike protein plasmids used in pseudovirus production. We are additionally grateful to Robert Kinsey, who conducted the manufacture of the NLC used in these studies; Christopher Press, who manufactured some RNA products used in this study; Jeff Guderian for molecular biology and laboratory managerial support; and Ethan Lo for immunoassay support. We also thank Theresa Britschgi, who was instrumental as project manager for this work, and Valerie Soza, for editorial assistance. We also gratefully acknowledge Thomas ‘Scott’ Simpler and Rebecca Burnham for ABSL2 hamster husbandry, Donovan J. Murphy and Robert Alldredge for ABSL3 hamster husbandry, Levi Schaefers and Sixto M. Leal Jr. (UAB Fungal Reference Lab) for viral RNA quantitation, Emily E. Helmen and Sherri Coffman (UAB Comparative Pathology Lab) for histology, and the UAB Immunology Institute’s Antibody Characterization and Serology (ACS) core for virus-specific antibody cytometric bead arrays.

## Author Contributions

Conceptualization: MFJ, EAV, FEL

Supervision: MFJ, EAV, AG, KSH, TDR, FEL

Project administration: MFJ, EAV, AG, DB, FEL

Funding acquisition: EAV, CC, FEL

Investigation: MFJ, SB, DH, PB, JB, DH, RM, JS, NC, SR, MDS, DB, AS-S, DNK, IRC, FZ, JLT, JA, GRK

Data curation: MFJ, MDS

Formal analysis: MFJ, SB, PB, NC, EAV, MDS

Writing – original draft: MFJ, EAV

Writing – review & editing: all authors

## Data and Materials Availability

The datasets generated and/or analyzed during the current study are available from the corresponding author on reasonable request. AAHI’s unique saRNA constructs and nanostructured lipid carriers are available for noncommercial research, academic non-clinical research, or other not-for-profit scholarly purposes undertaken at a non-profit or government institution, under the terms and conditions of an appropriate Material Transfer Agreement. Correspondence and material requests should be addressed to the corresponding author.

## Supplementary Materials

Figure S1. Spleen flow cytometry gating strategy.

Figure S2. Th1-Th2 balance of SARS-CoV-2 spike-responsive CD4^+^ cells after prime-boost immunization with SARS-CoV-2 saRNA-NLC vaccine.

Figure S3. Lung flow cytometry gating strategy.

Figure S4. Intranasal vaccination generates lung-resident SARS-CoV-2 spike-reactive T cells.

Figure S5. Vaccine antibody responses in naive cagemates of sham-vaccinated and vaccinated hamsters.

Figure S6. Correlation between body weight and serum anti-spike IgG titer in mice vaccinated with SARS-CoV-2 saRNA-NLC vaccine and experimentally infected with SARS-CoV-2.

Figure S7. Vaccination with SARS-CoV-2 saRNA-NLC vaccine reduces lung pathology in both vaccinated hamsters and naive cagemates irrespective of route of priming and boosting immunization.

Figure S8. Immune responses and viral load in vaccinated and infected hamsters and naïve cagemates.

