## Supplementary Information for "Intranasal self-amplifying RNA SARS-CoV-2 vaccine produces protective respiratory and systemic immunity and prevents viral transmission"

### Supplementary Figures

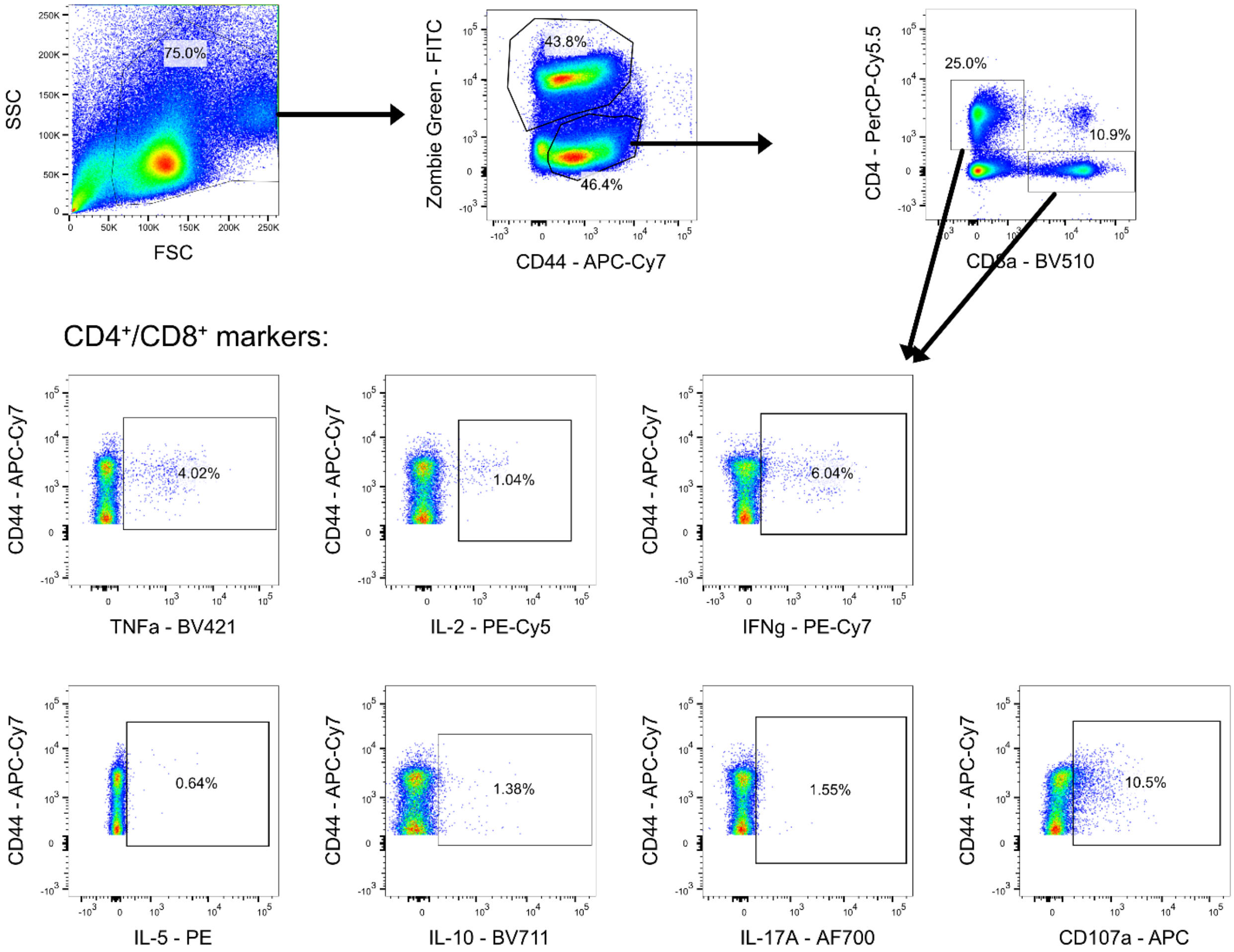


**Figure S1. Spleen flow cytometry gating strategy.** (Related to Figure 2). Isolated splenocytes were stained and analyzed via flow cytometry. Cells were gated first on total lymphocytes. The live cell subset was gated as FITC$^-$ and CD44$^+$. Then individual gates were drawn for CD4 (CD4$^+$CD8$^-$) and CD8 (CD4$^-$CD8$^+$) cells. Within each of these subsets, specific cytokines were gated using histograms, including IFNg, IL-2, TNFa, CD107a, IL-5, IL-10, and IL-17a.

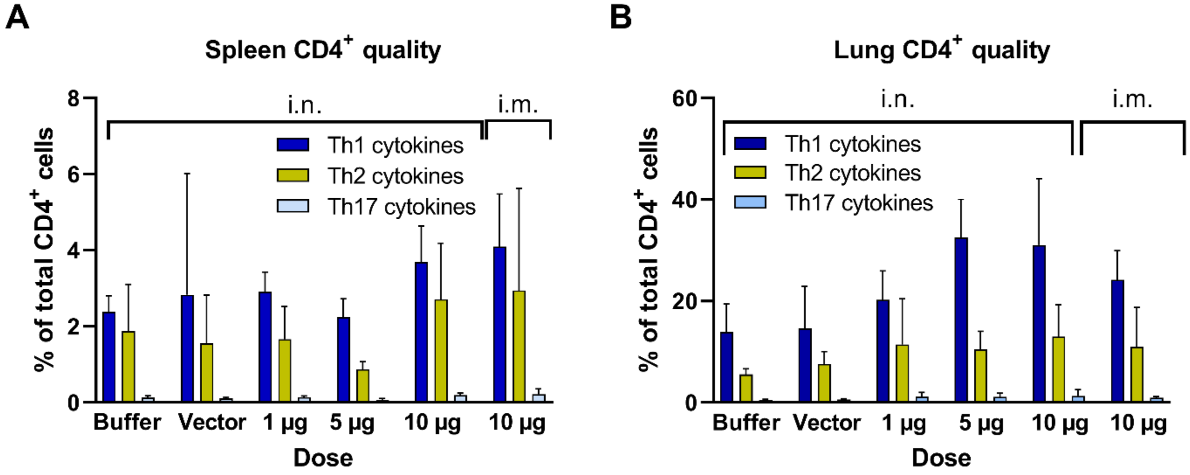


**Figure S2. Th1-Th2 balance of SARS-CoV-2 spike-responsive CD4$^+$ cells after prime-boost immunization with SARS-CoV-2 saRNA-NLC vaccine.** (Related to Figures 2 and 4.) **(A)** Quality of responding CD4$^+$ T cells. Total Th1, Th2, or Th17 responses indicated with bar graphs showing the average percentage across each group of mice ($n$ = 8 mice per group). **(B)** Quality of responding CD4$^+$ T cells, subdivided by cells that express one or more Th1 cytokine (IFNγ$^+$, IL-2$^+$, or TNFα$^+$), Th2 cytokine (IL-5), or Th17 cytokine (IL-17). $n$ = 8 mice per group. Vector control represents SEAP i.n. at 10 µg. Bars show mean ± SD. Statistics analyzed with one-way ANOVA with Tukey's multiple comparison test.

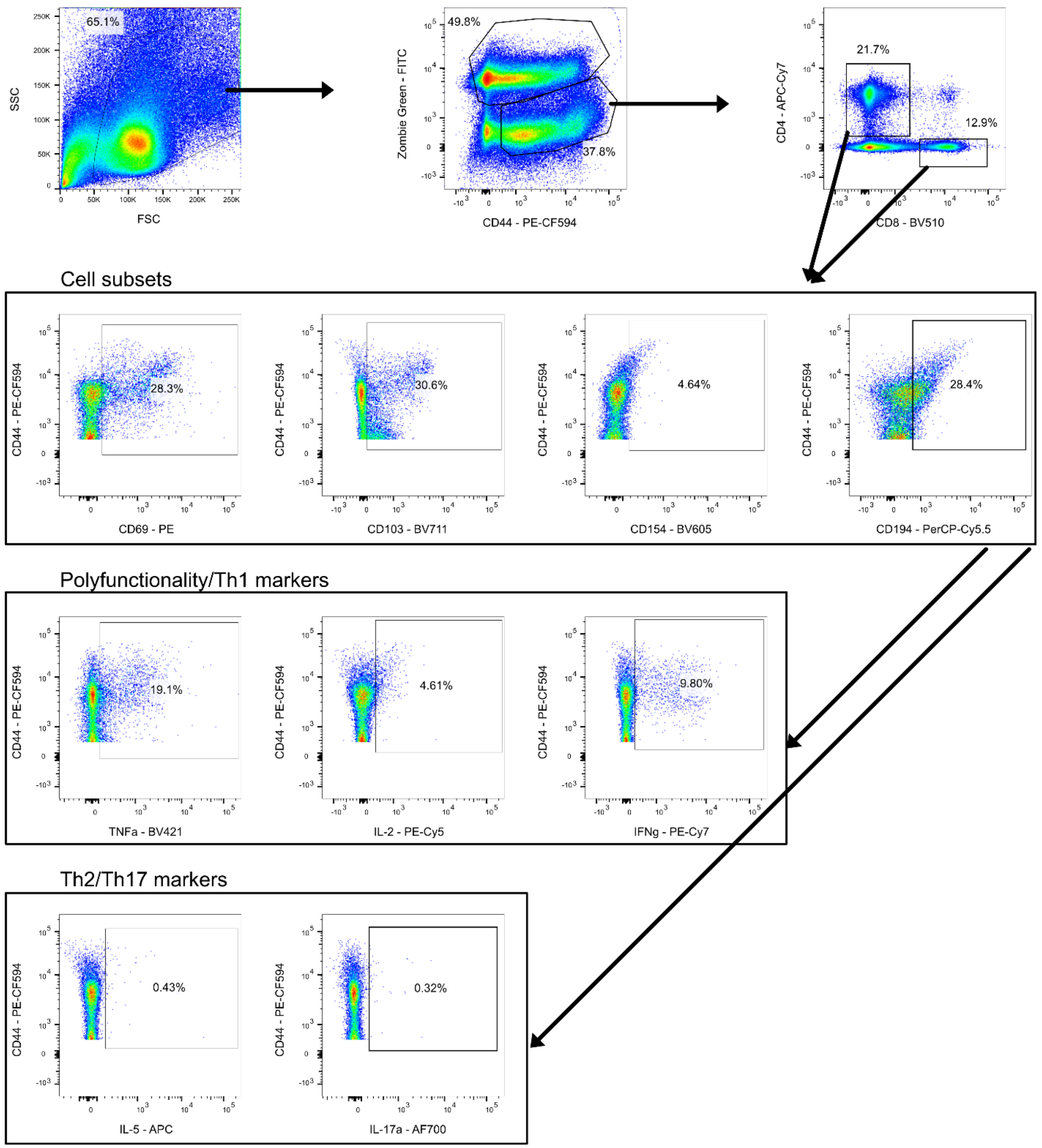


**Figure S3. Lung flow cytometry gating strategy.** (Related to Figure 3.) Isolated lung cells were stained and analyzed via flow cytometry. Cells were gated first on total lymphocytes. The live cell subset was gated as FITC$^-$ and CD44$^+$. Then individual gates were drawn for CD4 (CD4$^+$CD8$^-$) and CD8 (CD4$^-$CD8$^+$) cells. Within each of these subsets, specific cytokines were gated using histograms, including CD69, CD103 CD154, CD194, IFNg, IL-2, TNFa, IL-5, and IL-17a. Boolean gating was used to identify subsets of cells within other subsets.

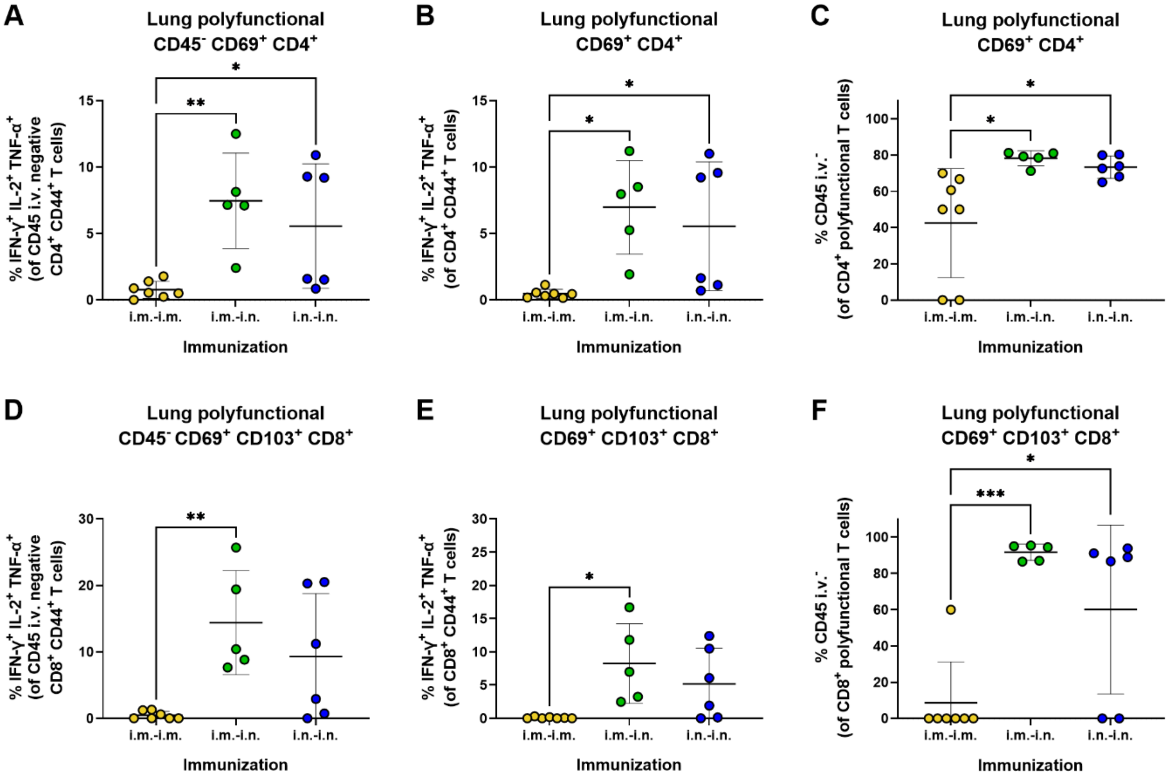


**Figure S4. Intranasal vaccination generates lung-resident SARS-CoV-2 spike-reactive T cells.** (Related to Figure 5). 6-8 week old C57BL/6J mice were immunized with 3 µg saRNA-NLC (N:P of 25) via i.m. or i.n. administration for prime and i.m or i.n. for boost dose 4 weeks later. **(A, D)** Frequency of lung CD45 intravenous (i.v.) label negative antigen-specific **(A)** $CD4^+$ and **(D)** $CD8^+$ T cells that are polyfunctional. **(B, E)** Frequency of lung antigen-specific polyfunctional **(B)** $CD4^+$ and **(E)** $CD8^+$ T cells. **(C, F)** Frequency of lung antigen-specific polyfunctional **(C)** $CD4^+$ and **(F)** $CD8^+$ T cells that are CD45 i.v. label negative. $n$ = 5-7 mice/group. Horizontal lines shown mean and SD. Significance was determined by one-way ANOVA with Tukey's multiple comparisons test: *$p < 0.05$, **$p < 0.01$, and *** $p < 0.001$.

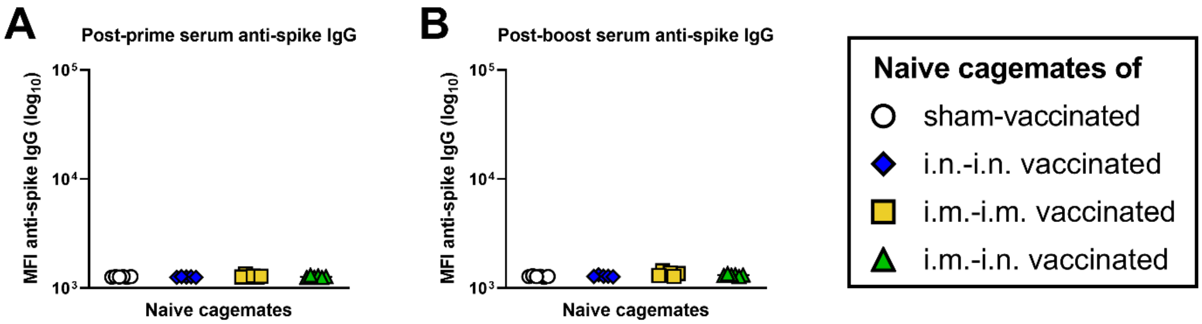


**Figure S5. Vaccine antibody responses in naive cagemates of sham-vaccinated and vaccinated hamsters.** (Related to Figure 6.) Vaccination regimens for vaccinated hamsters included sham-vaccinated (open black circles), i.n.-prime/i.n.-boost vaccinated (blue diamonds), i.m.-prime/i.m.-boost vaccinated (yellow squares), or i.m.-prime/i.n.-boost (green triangles). *n* = 5-6/group. Mean fluorescence intensity (MFI) of serum anti-spike IgG on **(A)** day 21 and **(B)** day 33. Horizontal lines show geometric mean.

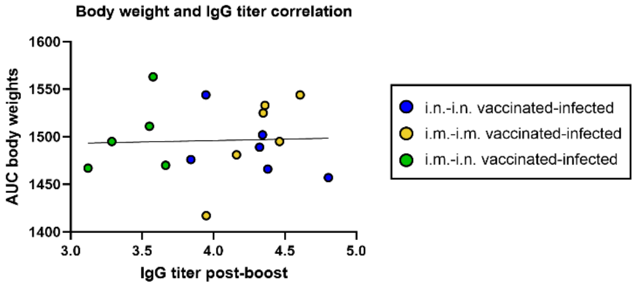


**Figure S6. Correlation between body weight and serum anti-spike IgG titer in mice vaccinated with SARS-CoV-2 saRNA-NLC vaccine and experimentally infected with SARS-CoV-2.** (Related to Figure 6). IgG titer in infected hamsters was correlated with the area under the curve (AUC) of hamster weight across the first 15 days of infection. IgG titer was log-transformed, and then Pearson correlation was calculated. Pearson $r^2$ =0.001497. $p$ = 0.8828. $n$ = 5-6 mice per group.

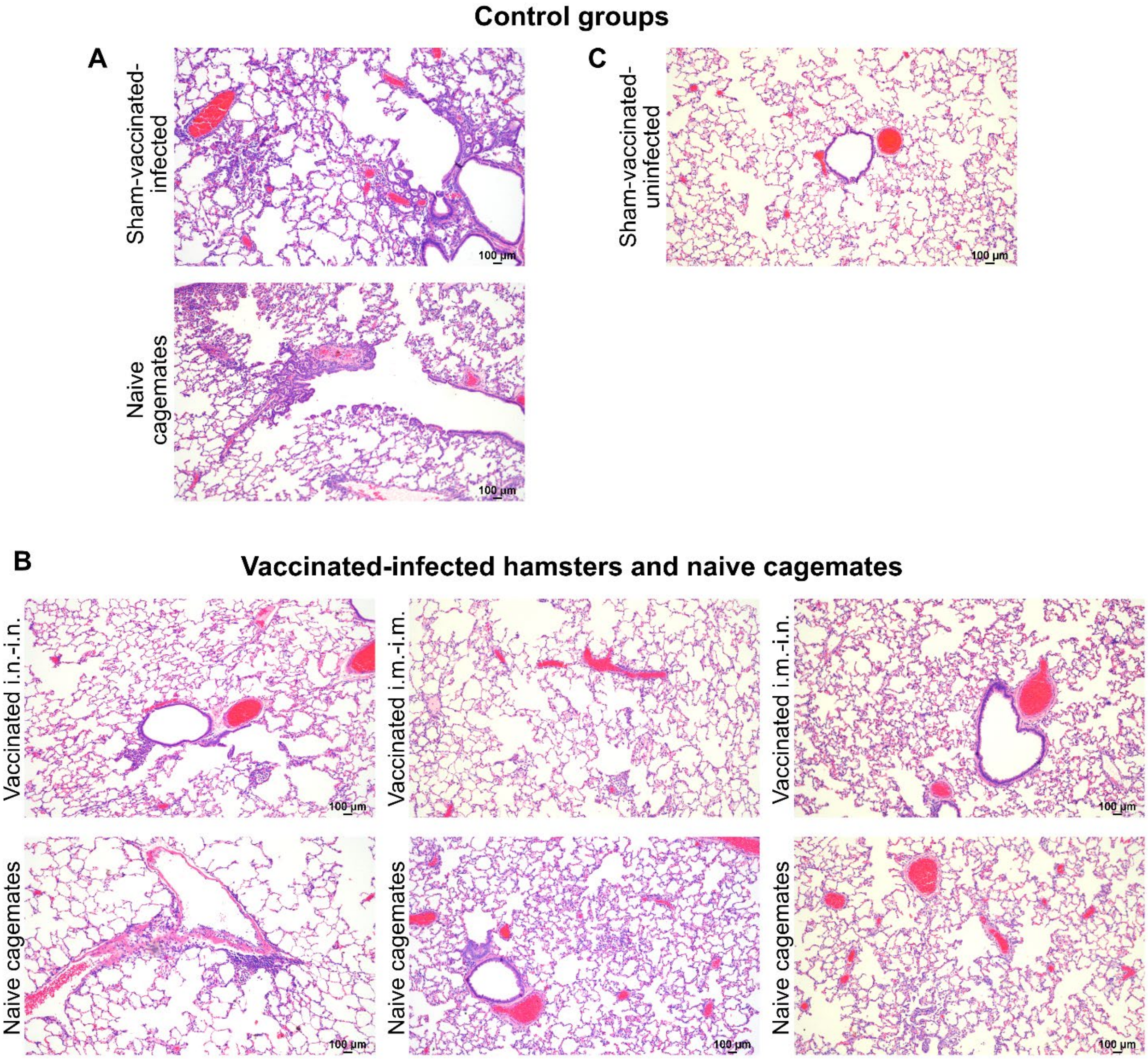


**Figure S7. Vaccination with SARS-CoV-2 saRNA-NLC vaccine reduces lung pathology in both vaccinated hamsters and naive cagemates irrespective of route of priming and boosting immunization.** (Related to Figure 6.) Hamsters from different vaccinated groups were infected with SARS-CoV-2 USA/WA-1 strain and, at 2 days post-infection, co-housed with naive cagemates. At 35 days post-infection, hamsters and their respective naive cagemates were euthanized, and lung tissue was evaluated for interstitial, perivascular, and peribronchiolar inflammation as represented in these 10X images. For comparison, sham-vaccinated-uninfected controls were assessed to determine presence of inflammation unrelated to SARS-CoV-2 infection. Control groups included **(A)** hamsters sham-vaccinated-infected or **(C)** sham-vaccinated-uninfected. Vaccinated groups included **(B)** hamsters vaccinated with an intranasal prime and boost (i.n.-i.n.), an intramuscular prime and boost (i.m.-i.m.), or an intramuscular prime and intranasal boost (i.m.-i.n.). Representative lung histology images from the naïve cagemates that were paired with animals from each of the groups are also shown.

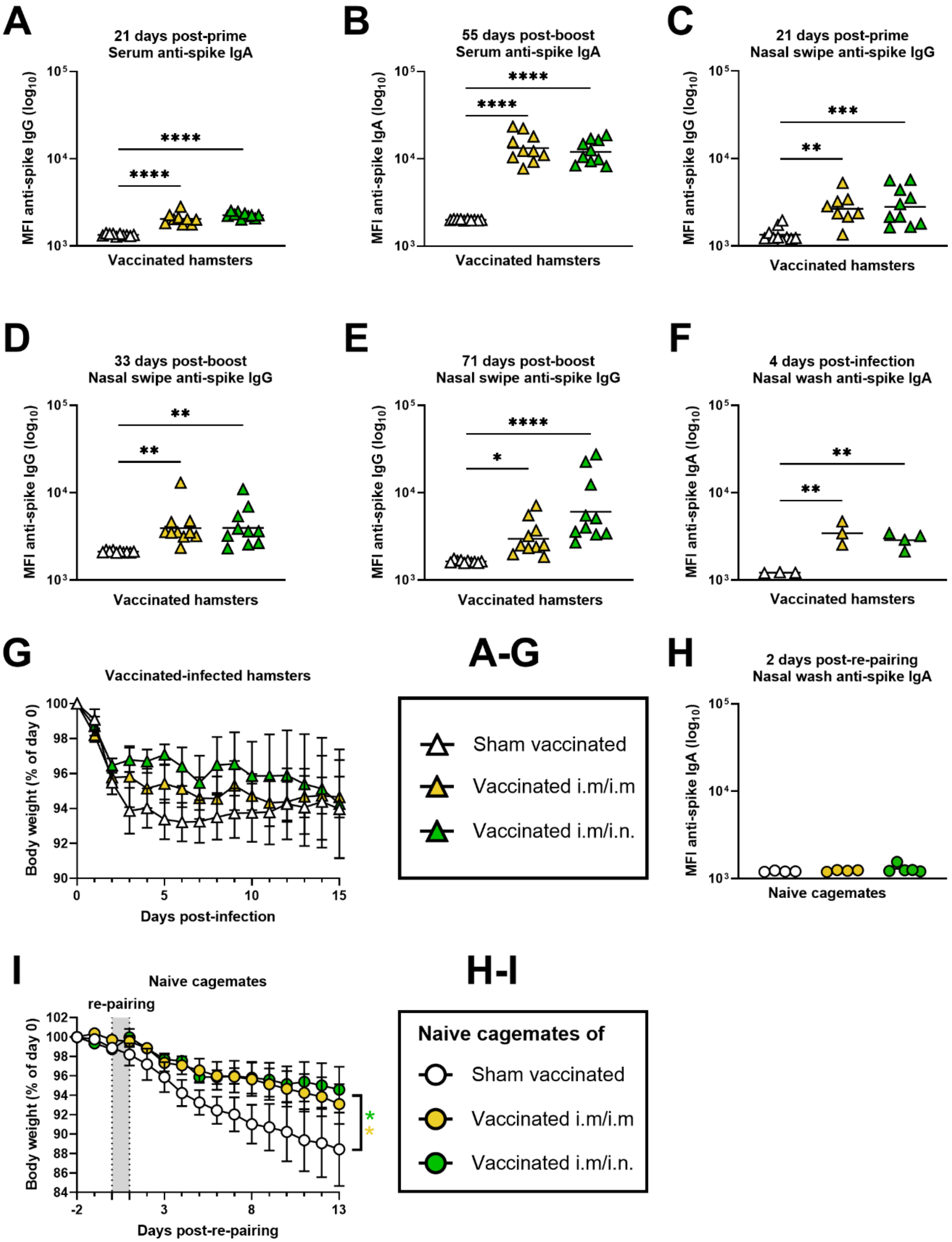


**Figure S8. Immune responses and viral load in vaccinated and infected hamsters and naïve cagemates.** (Related to Figure 7). **(A-F)** Vaccine-elicited antibody responses in sham-vaccinated (open white triangles), i.m.-prime/i.m.-boost vaccinated (yellow triangles), or i.m.-prime/i.n.-boost vaccinated (green triangles) animals ($n$ = 8-10/group). See Figure 7A for study design and outline. Anti-spike IgG and IgA levels reported as log-transformed MFI of antibody binding to SARS-CoV-2 spike-coated beads. The anti-spike IgA levels in serum on **(A)** day 21 and **(B)** 55 days post-boost (day 83), and the anti-spike IgG levels detected in nasal swabs on **(C)** day 21, **(D)** 33 days post-boost (day 61), and **(E)** 71 days post-boost (day 99) are reported. Data analyzed using one-way ANOVA with Tukey's multiple comparison test. Horizontal lines show geometric mean. (**F-G**) Immune responses in sham-vaccinated and prime+boost vaccinated hamsters infected 83 days following the boost with SARS-CoV-2 ($n$ = 4-5/group). **(F)** Spike-specific IgA antibodies in nasal wash isolated 4 days post-infection from the vaccinated-infected and sham-vaccinated-infected hamsters. Data reported as geometric mean of the log-transformed MFI values and analyzed using one-way ANOVA. **(G)** Body weight of vaccinated-infected and sham-vaccinated-infected hamsters. Data shown as group weight mean and SEM and analyzed as area under the curve using one-way ANOVA with Tukey's multiple comparison test. **(H-I)** Immune responses in naïve hamsters following re-pairing with sham-vaccinated-infected (open white circles), i.m.-prime/i.m.-boost vaccinated (yellow circles), or i.m.-prime/i.n.-boost vaccinated (green circles) cagemates ($n$ = 4-5/group). **(H)** Spike-specific IgA antibodies present in nasal wash of naive hamsters 2 days post-re-pairing with vaccinated-infected and sham-vaccinated-infected cagemates. Data shown as geometric mean and analyzed using one-way ANOVA. **(I)** Body weights of naive hamsters following re-pairing with prime+boost vaccinated-infected or sham-vaccinated-infected cagemates. Data shown as group weight mean and SEM. Data analyzed as area under the curve and assessed using one-way ANOVA with Tukey's multiple comparison test. * $p < 0.05$, ** $p < 0.01$, *** $p < 0.001$, **** $p < 0.0001$.
